# The UFMylation pathway is impaired in Alzheimer’s disease

**DOI:** 10.1101/2024.05.24.595755

**Authors:** Tingxiang Yan, Michael G. Heckman, Emily C. Craver, Chia-Chen Liu, Bailey D. Rawlinson, Xue Wang, Melissa E. Murray, Dennis W. Dickson, Nilufer Ertekin-Taner, Zhenkun Lou, Guojun Bu, Wolfdieter Springer, Fabienne C. Fiesel

**Affiliations:** Department of Neuroscience, Mayo Clinic, Jacksonville, FL 32224, USA; Division of Clinical Trials and Biostatistics, Mayo Clinic, Jacksonville, FL, USA; Neuroscience PhD Program, Mayo Clinic Graduate School of Biomedical Sciences, Jacksonville, FL 32224, USA; Department of Oncology, Mayo Clinic, Rochester, MN, 55905, USA

**Keywords:** UFMylation, UFM1, UFSP2, Alzheimer’s disease, tau, brain

## Abstract

**Background:** Alzheimer’s disease (AD) is characterized by the presence of neurofibrillary tangles made of hyperphosphorylated tau and senile plaques composed of beta-amyloid. These pathognomonic deposits have been implicated in the pathogenesis, although the molecular mechanisms and consequences remain undetermined. UFM1 is an important, but understudied ubiquitin-like protein that is covalently attached to substrates. This UFMylation has recently been identified as major modifier of tau aggregation upon seeding in experimental models. However, potential alterations of the UFM1 pathway in human AD brain have not been investigated yet.

**Methods:** Here we used frontal and temporal cortex samples from individuals with or without AD to measure the protein levels of the UFMylation pathway in human brain. We used multivariable regression analyses followed by Bonferroni correction for multiple testing to analyze associations of the UFMylation pathway with neuropathological characteristics, primary biochemical measurements of tau and additional biochemical markers from the same cases. We further studied associations of the UFMylation cascade with cellular stress pathways using Spearman correlations with bulk RNAseq expression data and functionally validated these interactions using gene-edited neurons that were generated by CRISPR-Cas9.

**Results:** Compared to controls, human AD brain had increased protein levels of UFM1. Our data further indicates that this increase mainly reflects conjugated UFM1 indicating hyperUFMylation in AD. UFMylation was strongly correlated with pathological tau in both AD-affected brain regions. In addition, we found that the levels of conjugated UFM1 were negatively correlated with soluble levels of the deUFMylation enzyme UFSP2. Functional analysis of UFM1 and/or UFSP2 knockout neurons revealed that the DNA damage response as well as the unfolded protein response are perturbed by changes in neuronal UFM1 signaling.

**Conclusions:** There are marked changes in the UFMylation pathway in human AD brain. These changes are significantly associated with pathological tau, supporting the idea that the UFMylation cascade might indeed act as a modifier of tau pathology in human brain. Our study further nominates UFSP2 as an attractive target to reduce the hyperUFMylation observed in AD brain but also underscores the critical need to identify risks and benefits of manipulating the UFMylation pathway as potential therapeutic avenue for AD.

## INTRODUCTION

Ubiquitin-fold modifier 1 (UFM1) is a small, ubiquitin-like protein that is covalently attached to lysine residues of substrate proteins in a process termed UFMylation[1, 2]. Similar to ubiquitylation, this post-translational modification is catalyzed by a series of enzymes. The first step is the maturation of the UFM1 precursor (proUFM1) by the UFM1-specific cysteine proteases (UFSP1 and UFSP2), which cleave the dipeptide Ser-Cys from the C-terminus to expose a single glycine residue that can be used for conjugation[3]. Subsequently, mature UFM1 is conjugated to target substrates via a catalytic cascade involving a UFM1-specific set of E1 (UFM1-activating enzyme - UBA5), E2 (UFM1-conjugating enzyme - UFC1), and a complex that consists of the E3 ligase (UFM1 ligase - UFL1) and the adaptor proteins DDRGK1 (aka UFBP1) and CDK5RAP3[4–8]. UFMylation is reversible. The deconjugation of UFM1 is mainly mediated by the protease UFSP2, loss of which significantly induces the accumulation of conjugated UFM1[9, 10].

The UFMylation pathway has been associated with a range of cellular functions, including unfolded protein response[11, 12], DNA damage response[13, 14], autophagic functions as well as immune response[15–20]. Interestingly, these cellular functions are central to neurodegeneration and Alzheimer’s disease (AD)[21–24]. Neuropathologically, AD is characterized by the presence of extracellular senile plaques composed of beta-amyloid (Aβ) and intracellular neurofibrillary tangles made of hyperphosphorylated forms of the microtubule-associated protein tau[25]. Very recently, a group identified UFMylation as novel key modifier of seeding-induced Tau propagation[26]. In addition, UFMylation is essential for brain development, as loss of function of any of its components causes severe neurodevelopmental disorders[15, 27–31]. Therefore, reduced functions of UFMylation could well affect neuronal function and viability. However, UFMylation and its role in and significance for neurodegenerative disorders are just emerging and changes in human AD brain have not been investigated yet.

In this study, we biochemically measured levels of UFMylation pathway proteins in temporal cortex and frontal cortex of control and AD brain. We assessed associations with primary clinical parameters and the severity of AD pathology, the abundance of AD-related molecules (tau, Aβ and APOE), as well as the expression of DNA damage and unfolded protein response related genes. This revealed a significantly increased abundance of the UFM1 protein in the cortex of AD brains, which was further associated with loss of soluble UFSP2 and the accumulation of pathological tau. Furthermore, we investigated the functional consequences of aberrant UFMylation in neurons and observed dual effects: protective benefits against DNA damage but increased susceptibility towards unfolded protein stress in neurons. Our study highlights disease-associated changes in UFMylation that might be associated with tau pathology in disease.

## MATERIALS AND METHODS

### Subjects

This study obtained de-identified post-mortem tissues from the Mayo Clinic Brain Bank. We analyzed two cohorts that each consisted of AD patients and of neurological normal individuals (hereafter referred to as controls). For the smaller exploratory cohort, we investigated frontal cortex samples from n=13 AD and n=13 controls. For the main cohort we investigated midfrontal and superior temporal cortex samples from n=72 AD and n=41 control cases. Detailed characteristics of these cohorts are summarized in **Tables S1 and S2**, respectively.

All brains were examined in a systematic and standardized manner and obtained between 1998 and 2019. All subjects are non-Hispanic Caucasians of European descent. Available clinical information included age at death, sex, Braak tangle stage (0-VI), and Thal amyloid phase (0–5) [33,62]. For the AD cohort we also obtained the age at onset, and disease duration. For the main cohort, we further obtained additional information such as the APOE genotype and mini mental state examination (MMSE) scores (AD patients only). The AD cases of the main cohort were part of the M²OVE-AD (Molecular Mechanisms of the Vascular Etiology of AD) initiative and had been phenotyped in depth. Levels of apoE, Aβ_40_, Aβ_42_, tau, pT231-tau were available from three fractions (Tris-buffered saline [TBS] buffer, detergent-containing buffer [1% Triton X-100 in TBS, termed TX], and formic acid [FA] fractions) from temporal cortex tissue[32]. These parameters were used as secondary measures of interest. In addition, we used bulk transcriptome data available from the same cases to study correlations with gene expression data.

The Mayo Clinic brain bank for neurodegenerative disorders operates with approval of the Mayo Clinic Institutional Review Board. All brain samples are from autopsies performed after approval by the legal next-of-kin. Research on de-identified postmortem brain tissue is considered exempt from human subjects regulations by the Mayo Clinic Institutional Review Board.

### Sample preparation

Cortex tissues were dissected and kept frozen until protein extraction. 180-200 mg of frozen tissue were homogenized in 5 volumes of ice-cold Tris-buffered saline (TBS; 50 mM Tris [Millipore, G48311], 150 mM NaCl [FisherScientific, BP358], pH 7.4) containing phosphatase inhibitors (Roche, 4906845001) and protease inhibitor cocktail (Roche, 11836170001) with a Dounce tissue grinder (DWK, K885300-0002). For protein extraction, ¼ volume of a 5x RIPA buffer (50 mM Tris, pH 8.0, 150 mM NaCl, 0.1% SDS, 0.5% deoxycholate, 1% NP-40) was added to the TBS homogenate and incubated at 4°C for 30 min with rotation. Then, samples were centrifuged at 100,000 g for 60 min at 4°C. The supernatant (referred to as ‘soluble’ fraction) was collected, aliquoted, flash frozen in liquid nitrogen and stored at -80°C until use. The residual pellet was washed with 1xRIPA buffer twice and centrifuged at 100,000 g for 30 min at 4°C. The pellet was resuspended in 2% SDS (Fisher, BP166-500) in TBS with phosphatase and protease inhibitors, sonicated for ten cycles (one cycle is 30 s ON/30 s OFF with high power level) in a Bioruptor plus sonication system (diagenode, Belgium) at 18 °C, and boiled at 95°C for 5 min. After centrifugation at 100,000 g for 60 min at 22°C, the resulting supernatant (referred to as ‘insoluble’ fraction) was collected, aliquoted, flash frozen in liquid nitrogen and stored at -80°C until use.

### Gel electrophoresis and western blot

The protein concentration was measured using BCA assay (Thermo Fisher, 23225). 20 μg protein extract was mixed with 6x SDS-PAGE loading buffer, boiled for 5 min at 95 °C and loaded on 8-16% Tris-Glycine gels (Invitrogen, EC60485BOX). Proteins were transferred onto 0.2 μm nitrocellulose membranes (Bio-Rad, 1620112). Following blocking with 5% nonfat milk (Sysco, 5398953) in TBS with 0.1% Tween (TBST) for one hour at room temperature (RT), primary and secondary antibodies were applied, and the blots developed with Immobilon Western Chemiluminescent HRP Substrate (Millipore, WBKLS0500). Bands were visualized on Blue Devil Lite X-ray films (Genesee Scientific, 30-810L) or with a ChemiDoc MP Imager (BioRad, Hercules, CA).

### Antibodies

The following antibodies were used for immunoblot: Rabbit anti-UFM1-Ab1 (Abcam, ab109305, 1:1000), rabbit anti-UFM1-Ab2 (Sigma, HPA039758, 1:1000), rabbit anti-UFM1-Ab3 (Proteintech Group, 15883-1-AP, 1:1000), rabbit anti-UFM1-Ab4 (LS Bio, LS-C807041, 1:1000), rabbit anti- UFM1-Ab5 (LS Bio, LS-C500000, 1:1000), mouse anti-UFSP1 (Santa Cruz Biotechnology, sc-398577, 1:1000), mouse anti-UFSP2 (Santa Cruz Biotechnology, SC-376084, 1:2000), rabbit anti-UBA5 (Proteintech Group, 12093-1-AP, 1:2000), rabbit anti-UFC1 (Abcam, ab189252, 1:2000), rabbit anti-UFL1 (Thermo Fisher, A303-456A, 1:1000), rabbit anti-DDRGK1 (Proteintech Group, 21445-1-AP, 1:1000), rabbit anti-CDK5RAP3 (Abcam, ab242399, 1:1000), mouse anti-β-actin (Sigma, A1978, 1:100,000), mouse anti-Vinculin (Sigma, V9131, 1:100,000), mouse anti-GAPDH (Meridian Life science, H86504M; 1:5,000,000), rabbit anti-Bip (Cell Signaling Technology, 3177, 1:5,000), rabbit anti-PERK (Cell Signaling Technology, 3192, 1:4000), rabbit anti-ATF4 (Cell Signaling Technology, 11815, 1:2000), mouse anti-CHOP (Cell Signaling Technology, 2895, 1:2000), rabbit anti-IRE1α (Cell Signaling Technology, 3294, 1:2000), rabbit anti-Xbp1s (Cell Signaling Technology, 12782, 1:5000), rabbit anti-ATF6 (Cell Signaling Technology, 65880, 1:1000).

The following antibodies were used for immunofluorescence: mouse anti-CHOP (Cell Signaling Technology, 2895, 1:200), rabbit anti-Xbp1s (Cell Signaling Technology, 12782, 1:400), γH2Ax (Cell Signaling Technology, 9718T, 1:400).

For ELISA the following antibodies were used: rabbit anti-UFM1-Ab1 (Abcam, ab109305, 1:300), rabbit anti-UFM1-Ab2 (Sigma, HPA039758, 1:100), rabbit anti-UFM1-Ab3 (Proteintech Group, 15883-1-AP, 1:100), rabbit anti-UFM1-Ab4 (LS Bio, LS-C500000, 1:100), rabbit anti-UFM1-Ab5 (LS Bio, LS-C807041, 1:100), mouse anti-UFSP2 (Santa Cruz Biotechnology, SC-376084, 1:50), rabbit anti-tau (DAKO, AA002402-1, 1:500), mouse anti-total tau (Invitrogen, AHB0042, 1:500), mouse anti-p-tau (PHF1, a generous gift from the late Dr. Peter Davies, 1:500).

### Generation of gene-edited neuronal precursor cells

Neuronal progenitor cells derived from the ventral mesencephalon (ReN cell VM, Millipore, SCC008) were maintained on growth factor-reduced matrigel (Corning, CB-40230) coated plates in DMEM-F12 media (Thermo Fisher, 11320033), supplemented with B27 (Thermo Fisher, 17504044), 50 µg/ml gentamicin (Thermo Fisher, 15-750-060), and 5 U/ml Heparin (Sigma, H3149) in the presence of 20 ng/ml epidermal growth factor (EGF, Peprotech, AF-100-15) and fibroblasts growth factor (FGF, Peprotech, 100-25). Differentiation of ReN cells was performed by replacing FGF and EGF with 2 ng/ml GDNF (Peprotech, 450–10) and 1 mM dibutyryl-cAMP (Invivochem, V1846) for fourteen days[33]. All cells were grown at 37°C, 5% CO2/air in a humidified atmosphere.

We used ALT-R CRISPR-Cas9 system (IDT, Coralville, IA) to knock out UFM1 or UFSP2. UFM1 was further knocked out in UFSP2 KO cells to generate double knockouts (dKO). ReN cells VM were electroporated with ribonucleoprotein complex using the nucleofector P3 kit (Lonza, V4XP-3032). Single cell colonies were generated by limited dilution in 96-well plates. All clones were analyzed by PCR and western blot. The five most likely off-target sites as identified by the Benchling biology software (2021, www.benchling.com) were sequenced by Sanger sequencing to exclude unwanted editing. The sequences of gRNAs were as follows: gRNA-UFM1: GTAAGCAAACACTTACATGG; gRNA-UFSP2: AATAAGAGGAGGCCTTGATT.

### Quantification of UFM1, UFSP2, total tau, and pS396/404-tau

The relative amounts of UFM1, UFSP2, tau and pS396/404-tau were measured by Meso Scale Discovery (MSD) ELISA. All samples were run in duplicates. For the UFM1 and UFSP2 MSD ELISA, 10 μg of denatured brain samples were diluted in 200 mM sodium carbonate buffer pH 9.7 overnight at 4°C in 96-well MSD plates (MSD, L15XA-3). Plates were washed 3 times with 300 μl TBST wash buffer, blocked with 5% nonfat milk in TBST for one hour at RT, then incubated with primary antibody for UFM1 (Abcam, ab109305) or UFSP2 (Santa Cruz Biotechnology, sc-376084) diluted in 5% nonfat milk for 2 hours at RT using agitation, washed 3 times with TBST, and incubated with SULFO-TAG labeled goat anti-rabbit (for UFM1, MSD, R32AB-1) or anti-mouse (for UFSP2, MSD, R32AC-1) for 1 h at RT using agitation. After the final three washing steps, 150 μl MSD GOLD Read Buffer (MSD, R92TG-2) was added to each well and the plate read on a MESO QuickPlex SQ 120 reader (MSD, Rockville, MD, USA). Lysates of UFM1 or UFSP2 KO ReN cells were used as negative controls.

Levels of total tau were determined by the sandwich MSD ELISA using a polyclonal total tau antibody (DAKO, A0024) as a capture antibody and a monoclonal total tau antibody (TAU-5, Thermo, AHB0042) as a detection antibody. Levels of phosphorylated (pS396/404) tau were determined by a MSD sandwich ELISA using a polyclonal total tau antibody (DAKO, AA002402-1) as a capture antibody and a monoclonal pS396/404-tau antibody (PHF1) as a detection antibody.

### Cell treatments, staining and microscopy

Neuronal progenitor cells were plated on matrigel coated 96-well plates (PerkinElmer, 6055302) and differentiated for 14 days. DNA damage was induced with 10 µM etoposide (Cayman Chemical, 12092-25) for analysis of γH2Ax immunostaining and with 100 µM etoposide or 10 µM bleomycin (Sigma, B1141000) for cell viability analysis. ER stress was induced by treatment of cells with 10 µg/ml tunicamycin (Sigma, T7765) or 1 µM thapsigargin (Santa Cruz Biotechnology, sc-24017).

For immunostaining, cells were fixed with 4% paraformaldehyde (Thermo Scientific Chemicals, J19943.K2) for 10 minutes, washed with PBS (Boston Bioproducts, BM-220) three times before permeabilization with 0.1% Triton X-100 in PBS for 10 min at RT. After blocking with 10% normal goat serum (Invitrogen, 16210072) in PBS, cells were stained with γH2AX (Cell Signaling Technologies, 9718, 1:400), or Xbp1s (Cell Signaling Technologies, 40435, 1:400), and CHOP (Cell Signaling Technologies, 2895, 1:200) antibodies for 1.5 h, followed by secondary antibodies (donkey anti-rabbit IgG Alexa Fluor 488, donkey anti-mouse IgG Alexa Fluor 568, Thermo Fisher Scientific, A21206, A10037) for 1 h at RT. Nuclei were counterstained with Hoechst 33342 (1:5000 in PBS). For cell viability staining, a LIVE/DEAD Assay Kit (Invitrogen, L32250) was used according to the manufacturer’s instructions.

Imaging plates were imaged on an Operetta CLS system (PerkinElmer, Waltham, MA) with a 20x water objective using at least 4 fields per view per well (no gaps). Raw images were processed using the built-in Harmony software (version 4.9). Nuclei were identified based on the Hoechst staining and defined as regions of interest using the standard analysis building block. The mean fluorescence intensity of γH2AX, Xbp1s or CHOP was recorded for each nucleus and averaged. At least 1000 cells per genotype and condition were measured per experiment. Live cells were identified by a linear classifier that was developed using the integrated Phenologic machine learning module trained with intensity data for the live dye.

### Statistical analysis

Continuous variables were summarized with the sample median and range. Categorical variables were summarized with number and percentage. Comparisons of subject characteristics between AD patients and controls were made using a Wilcoxon rank sum test (continuous and ordinal variables) or Fisher’s exact test (categorical variables). Unadjusted pair-wise correlations between variables were assessed using Spearman’s test of correlation; p values below 0.05 were considered statistically significant in these exploratory analyses.

Comparisons of UFSP2 and UFM1 between AD patients and controls were made using unadjusted and age/sex-adjusted linear regression models. Soluble UFSP2 was examined on the square root scale in all analyses owing to its skewed distribution. Regression coefficients (denoted as β) and 95% confidence intervals (CIs) were estimated and are interpreted as the increase in mean UFSP2, or UFM1 (on the square root scale for soluble UFSP2) for AD cases compared to controls. In order to adjust for multiple testing for the primary comparisons of UFSP2 and UFM1 between AD patients and controls, we utilized a Bonferroni correction separately for the temporal and frontal cortices and separately for each outcome, after which p-values <0.025 were considered as statistically significant.

In the separate groups of controls and AD patients, associations of UFSP2 and UFM1 with clinical and disease parameters were evaluated using unadjusted and multivariable linear regression models, Multivariable models for controls were adjusted for age, sex, Braak stage, and Thal phase, while multivariable models for AD patients were adjusted for age, sex, presence of *APOE* ε4, Braak stage, and Thal phase. β coefficients and 95% CIs were estimated and are interpreted as the increase in mean UFSP2 (on the square root scale when examining soluble UFSP2) corresponding to presence of the given characteristic (categorical variables) or a specified increase (continuous variables). Continuous variables were examined on the untransformed, square root, cube root, or natural logarithm scale in regression analysis (**Table S3**). In order to examine associations of UFSP2 and UFM1 with clinical and disease parameters in the overall group of all subjects, we combined results for the separate AD and control groups using a random-effects meta-analysis[34]. We adjusted for multiple testing as follows: For the association analysis assessing correlations of UFM1 and UFSP2 with each other as well as with age, sex, *APOE* ε4, Braak stage, Thal phase, pS396/404-tau, and total tau, we applied a Bonferroni correction for multiple testing separately for each patient group, cortex, and fraction, after which p-values <0.01 (controls and all subjects) and <0.0071 (AD patients) were considered as statistically significant.

All statistical tests were two-sided. Spearman’s analysis and Wilcoxon rank sum tests were performed using GraphPad Prism (version 10.0.0, Boston, MA, USA). All other statistical analysis was performed using R Statistical Software (version 4.0.3; R Foundation for Statistical Computing, Vienna, Austria).

## RESULTS

### UFMylation pathway genes are differentially expressed in excitatory neurons of AD patients

To shed light onto the role of UFMylation for AD, we first performed a meta-analysis of published single nuclei transcriptome data from brain of patients with AD and controls (no-AD pathology)[35]. We compared expression levels of all UFMylation pathway components (**Fig. 1A**) across cell types including excitatory neurons, inhibitory neurons, astrocytes, oligodendrocytes, oligodendrocyte precursor cells, and microglia. Five of the eight UFMylation components (UFSP1, UFSP2, UFC1, UFL1, and DDRGK1) were significantly decreased in the excitatory neurons of AD brains (**Fig S1A, Table S4**). Other cell types showed either no or a lower differential expression of UFMylation genes between normal and AD brain (**Fig S1A, Table S4**). Of note, other ubiquitin-like pathways, such as ISGylation, NEDDylation, SUMOylation, or others did not show a comparable change (**Fig S1B, Table S4**), suggesting that the UFMylation pathway might be specifically altered in AD excitatory neurons.

**Figure 1:**
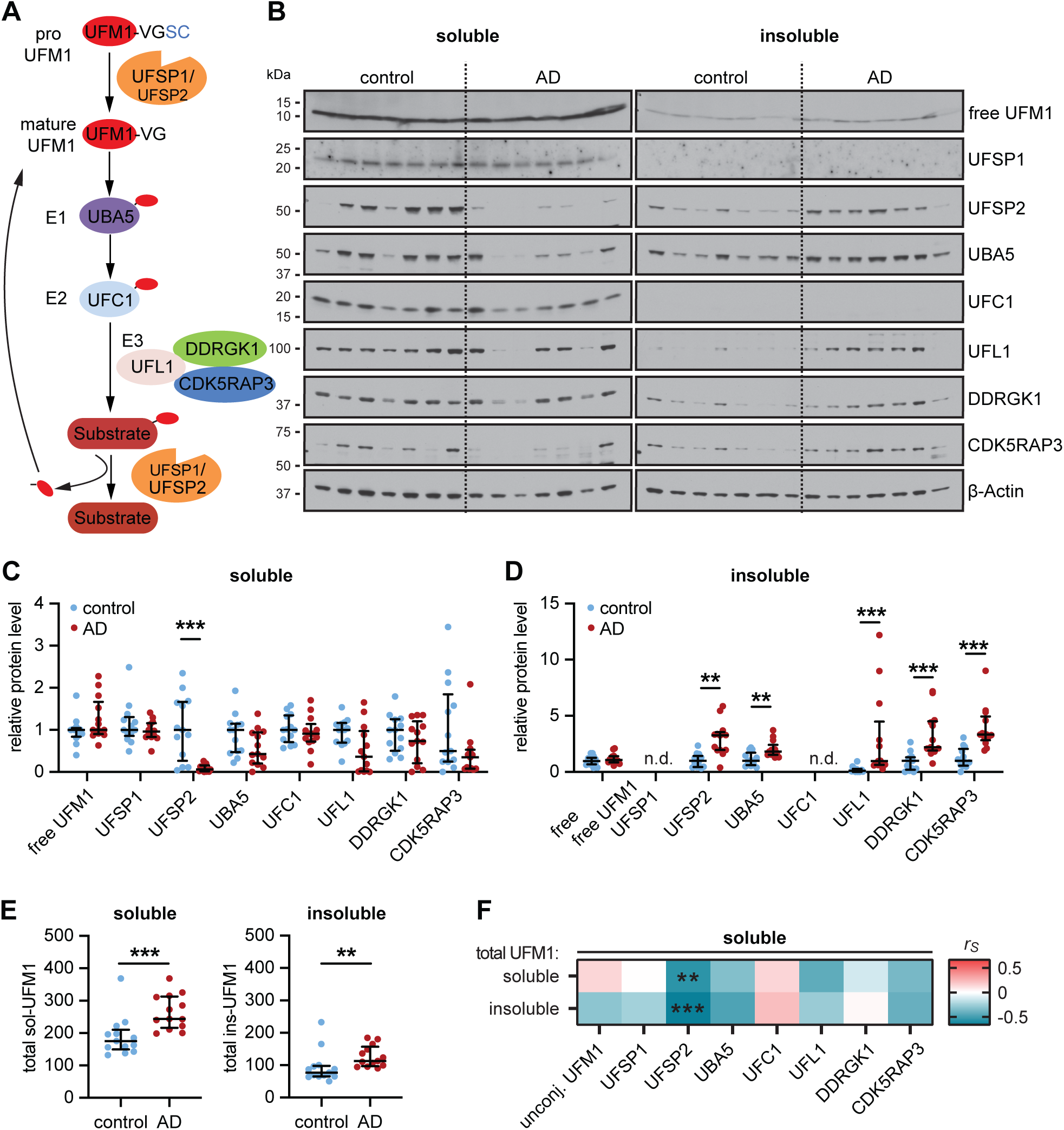
Exploratory analysis of the UFM1 pathway in normal and AD frontal cortex. (**A**) Schematic of the UFMylation pathway: Pro-UFM1 is cleaved by the protease UFSP1 or UFSP2 into the mature, conjugatable form. UBA5 (E1) activates UFM1 and UFC1 acts as an E2 conjugating enzyme that interacts with the E3 complex consisting of UFL1 and the adaptor proteins DDRGK1 and CDK5RAP3, which mediate the transfer of UFM1 from UFC1 to its target substrate. UFM1 is cleaved from its substrates mainly by UFSP2. (**B**-**D**) Representative immunoblot (B) and densitometric quantification of UFM1 pathway proteins in radioimmunoprecipitation assay (RIPA) buffer soluble (C) and insoluble (D) fractions of human normal and AD frontal cortex. UFM1 pathway protein levels were normalized to loading control beta-Actin and normalized to the median of the control cohort. Statistical analysis was performed using a Wilcoxon rank sum test followed by Bonferroni correction for testing two fractions, **P<0.00625, ***P<0.001; n.d. - not detected. (**E**) Quantification of total UFM1 via Meso Scale Discovery (MSD) enzyme-linked immunosorbent assay (ELISA). Data is shown as median with interquartile range. Statistical analysis was performed with Wilcoxon rank sum test followed by adjustment with Bonferroni correction for analyzing two fractions, *P<0.025, **P<0.01, ***P<0.001. (**F**) Soluble UFSP2 western blot levels are negatively correlated with soluble and insoluble total UFM1 levels that were determined by MSD ELISA. Shown is a heatmap of Spearman correlation coefficients (r_S_), **P<0.00625, ***P<0.001. n = 13 per group. See Supplementary Table S5 for r_s_ and p values.

### UFM1 and UFSP2 are altered in the frontal cortex of an exploratory AD cohort

To examine whether also protein levels of UFMylation pathway components are altered in human AD brain, we first used an exploratory cohort consisting of frontal cortex samples from 13 neurologically normals (hereafter to referred to as controls) and 13 AD subjects (see Table S1). All eight UFMylation pathway components were analyzed by western blot in the RIPA-soluble and -insoluble fraction (**Fig 1B-D**). Free, unconjugated UFM1 was not altered between AD and controls. However, protein levels of UFSP2 were significantly decreased in the soluble fraction (**Fig 1C**), while concurrently increased in the insoluble fraction of AD cases. In line with a general increase of aggregated proteins in AD, several other UFMylation proteins (UBA5, UFL1, DDRGK1 and CDK5RAP3) were also significantly increased in AD versus controls (**Fig 1D**). However, in contrast to UFSP2 the soluble portion of these other UFMylation proteins remained unchanged (**Fig 1B,C**).

UFPS2 is one of the UFM1-specific proteases. While UFSP1 and 2 have been both described with to facilitate pre-processing and recycling of UFM1[3], it is becoming increasingly clear that UFSP1 might be primarily important for the maturation of UFM1, while UFSP2 is important to cleave off UFM1 from its substrates[9, 10]. A loss of soluble UFSP2 in brain could therefore be linked to an increase of substrate-conjugated UFM1. Because UFM1 is attached to different substrates with distinct molecular weights, conjugated UFM1 appears as multiple bands or a smear in western blot, similar to ubiquitin (**Fig S2A**). However, none of the tested UFM1 antibodies was fully specific (**Fig S2B**) as some bands were still visible in samples from UFM1 knock out (KO) cells. To overcome these limitations, we developed a new Meso Scale Discovery (MSD) ELISA method (**Fig S2C,D**). With this assay, signal obtained with lysates from UFM1 KO was as low as the background signal without lysate added (buffer blank). In addition, lysates from UFSP2 KO neurons, which indeed show more conjugated UFM1 on western blot (**Fig S2B**), resulted in a higher signal compared to isogenic wild-type (WT) neurons, confirming that the assay detects total (i.e. conjugated and unconjugated) UFM1.

Using this UFM1 MSD ELISA on post-mortem brain samples, we found that total UFM1 levels were increased in both the soluble and insoluble fraction in AD compared to controls (**Fig 1E**). Since levels of unconjugated UFM1 were similar between AD and controls, the increase in total UFM1 likely reflects primarily an increase in conjugated UFM1. A correlation analysis between UFM1 levels, as determined by ELISA, and all other UFMylation pathway components that were determined by western blot revealed a significant negative correlation of soluble UFSP2 with both soluble (P=0.0028) and insoluble total UFM1 (P=0.0002) (**Fig 1F, Table S5**). To measure the protein level of UFSP2 on a larger scale, we developed another MSD ELISA that we validated with UFSP2 KO cells (**Fig S3A,B**). The UFSP2 levels obtained with this MSD ELISA correlated highly with levels assessed by western blot of AD and control samples (r=0.93, P=6.6x10^-12^) (**Fig S3C**). Consistently, ELISA-measured UFSP2 levels were also significantly different between controls and AD (**Fig S3D**). Furthermore, there was significant negative correlation with both soluble (P=0.0041) and insoluble total UFM1 (P=0.0029) (**Fig S3E**). Given the significant changes of total UFM1 and UFSP2 in our exploratory cohort, we decided to focus on these two UFMylation pathway members for further investigation.

### Expression of UFM1 and UFSP2 are altered in the temporal and frontal cortex in AD

We next studied a much larger cohort that consisted of 41 normal controls and 72 AD cases with similar sex and age (**Table S2**). The superior temporal cortex and the frontal cortex were included as early or later affected brain region, respectively. The AD cases of this cohort have previously been deeply phenotyped by biochemistry and using bulk RNAseq[32]. As expected, and consistent with the selection of subjects, both Braak tangle stage and Thal amyloid phase were significantly higher in AD cases compared to controls (see Table S2).

Consistent with findings from the pilot cohort, protein levels of both soluble and insoluble UFM1, as well as insoluble UFSP2, were all significantly increased in both brain regions from the larger AD group compared to controls as measured by MSD ELISA. Although soluble UFSP2 was only significantly decreased in the temporal cortex of AD patients, a trend was also noticeable in the frontal cortex that is later affected in disease (**Fig 2A**). In multivariable analysis adjusting for age and sex, compared to controls, there were significantly (P<0.025 considered significant) higher levels of soluble UFM1 in the temporal cortex (P=0.017), higher levels of insoluble UFSP2 in the frontal cortex (P=0.017), as well as higher levels of both soluble (P=0.002) and insoluble UFM1 (P<0.001) in the frontal cortex of AD patients (**Table 1**). Though not quite significant, there were trends towards higher insoluble UFSP2 levels in the temporal cortex of AD patients compared to controls (P=0.050), and towards higher levels of insoluble UFM1 in the temporal cortex of AD patients also approached significance (P=0.062) (**Table 1**).

**Figure 2:**
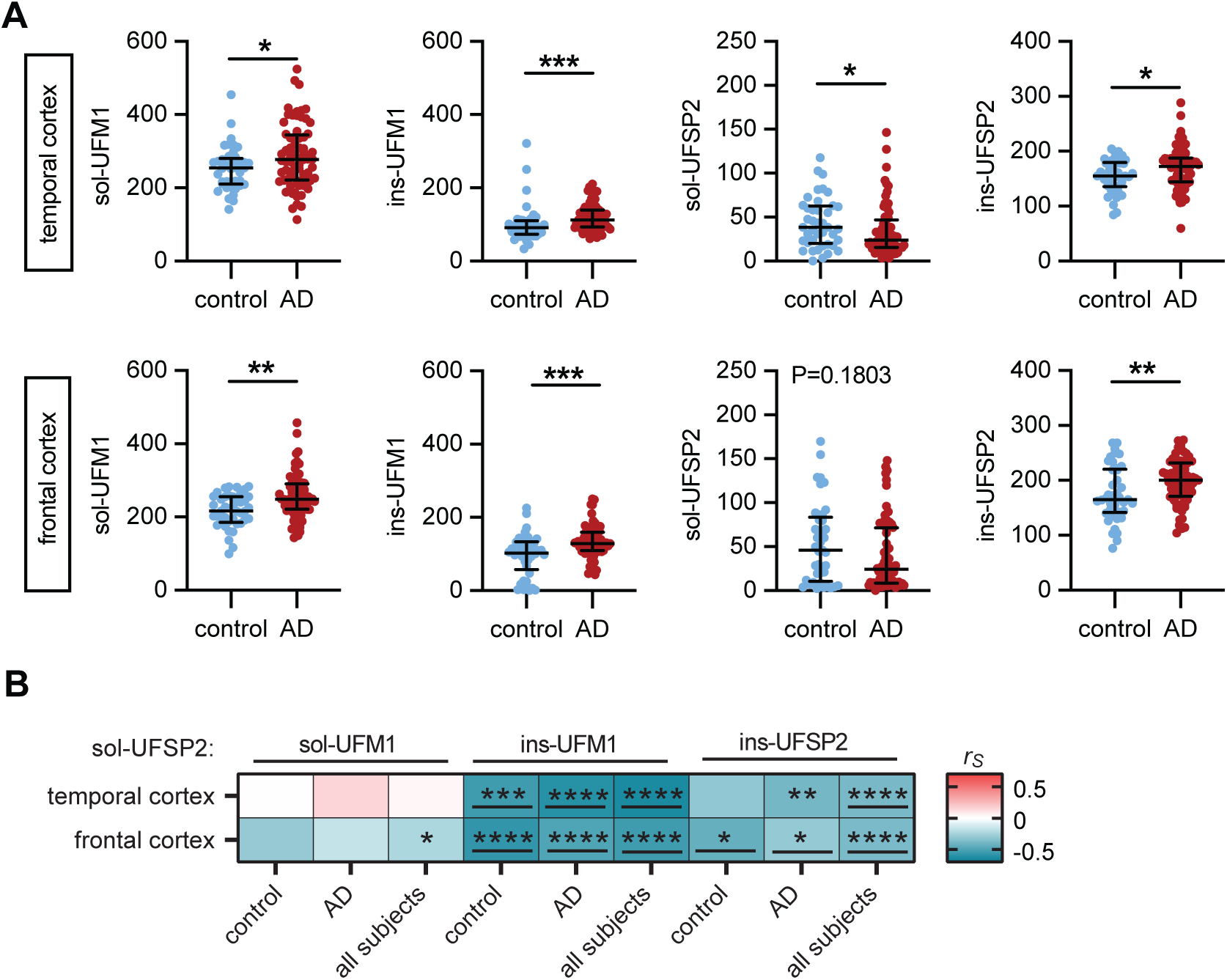
UFM1 and UFSP2 levels are altered in human AD brain. (**A**) Quantification of RIPA-soluble (sol) and -insoluble (ins) UFM1 and UFSP2 levels, respectively, by MSD ELISA in the frontal and temporal cortex of AD cases (n=72) and controls (n=41). Median and interquartile range is indicated. Statistical analysis was performed with a Wilcoxon rank sum test, *P < 0.05, **P<0.01, ***P<0.001. Linear regression analysis is summarized in Table 1. **(B)** Heatmap of Spearman correlation coefficients (r_s_) illustrating strong correlation between soluble UFSP2 with mostly insoluble UFM1 and UFSP2 levels from temporal cortex or frontal cortex of controls, AD and of combined cases (control + AD, n=113). Indicated significance levels are from Spearman’s test after Bonferroni correction: *P<0. 0167, **P<0. 0033, ***P<0. 0003, ****P<0.0001. Significant correlations that were confirmed by multivariable linear regression analysis (Supplementary Table S6) have been underlined.

**Table 1:**
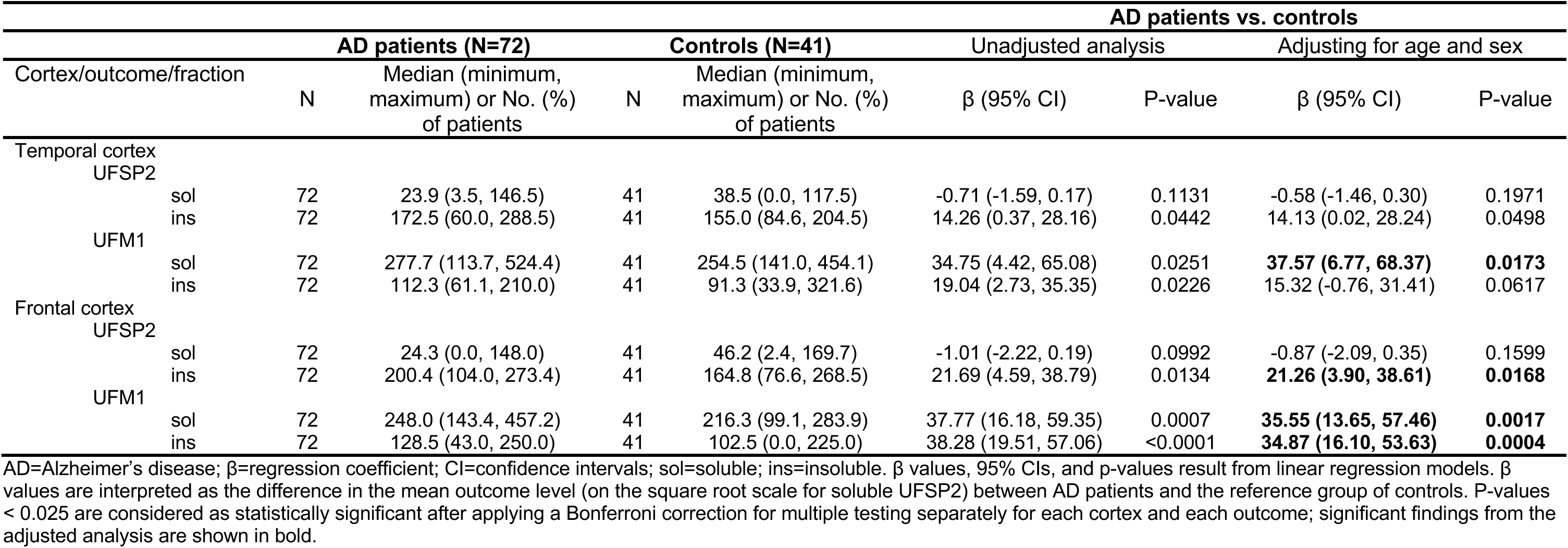
Comparisons of primary outcomes between AD patients and controls.

The negative correlation between UFSP2 and UFM1 that was observed in the pilot cohort was overall conserved (**Fig 2B**). However, in this larger cohort the correlation was mostly restricted to insoluble UFM1. Significant correlation with soluble UFM1 was only observed in the frontal cortex when all subjects were combined. The correlation of UFSP2 with insoluble UFM1 was stronger, and present in both brain regions and in all three groups (controls, AD, and when combining all subjects), and remained significant in multivariable linear regression models and upon adjusting for multiple testing (**Fig 2B, Table S6**). These results suggest that reduction of soluble UFSP2 levels may be associated with UFM1 accumulation, particularly in the insoluble fraction of the AD group in both the temporal and frontal cortices. Moreover, similar to the pilot cohort, there was a significant negative correlation between soluble and insoluble UFSP2 in both brain regions, pointing towards a solubility shift of UFSP2 (both P<0.001) (**Fig 2B, Table S6**).

### Levels of UFM1 are associated with pathological tau in human brain

To investigate whether the levels of UFM1 and UFSP2 are associated with primary clinical parameters and the severity of AD pathology, we first performed association analyses. After adjusting for multiple testing, there were only two significant associations between UFM1 or UFSP2 with clinical or pathological parameters such as age at death, sex, presence of *APOE* ε4 allele, disease duration, MMSE scores, Braak neurofibrillary tangle stage, and Thal amyloid phase (**Tables S7**). Specifically, levels of insoluble UFSP2 were significantly associated with Braak stage in the frontal cortex of the control, but not the combined or the AD cohort, while levels of insoluble UFM1 were associated with the Thal stage in the group that contains data from all subjects.

To explore the association between the levels of UFM1 and UFSP2 and pathological AD markers, we next obtained published data from the biochemical quantification of apoE, Aβ40, Aβ42, tau, and pT231-tau of sequential fractions of the temporal cortex from the same AD cases[32]. There was a significant correlation between soluble UFM1 and tau-related proteins, specifically total and pT231-tau in the TX fraction. Similarly, insoluble UFM1 also demonstrated significant correlations with pT231-tau in the TX and FA fraction (**Fig S5, Table S8**). In contrast, no significant correlations were established between the levels of apoE, Aβ40 and Aβ42 proteins and those of UFM1 and UFSP2 in the temporal cortex of AD patients. These findings highlight a closer association of UFM1 and UFSP2 levels with tau over other AD-related markers.

In order to investigate the relationship with tau further, we measured total tau and pS396/404-tau level in both the controls and AD cases with the MSD ELISA. We chose to focus on pS396/404-tau because it is associated with advanced stages of AD, unlike pT231-Tau, which is linked to early tau pathology changes[36]. Consistently, levels of pathological tau (soluble and insoluble pS396/404-tau, and insoluble total tau) were significantly higher in AD than controls in both brain regions (all P<0.0001, **Table S9**), while the levels of soluble total tau did not differ noticeably between these two groups. Higher levels of soluble UFSP2 were correlated with higher soluble total tau (P<0.001) in both cortices, indicative of association with physiological tau (**Fig. 3**). In contrast insoluble UFSP2, as well as soluble and insoluble UFM1 were correlated with pathological forms of tau (insoluble total tau and pS396/404-tau). These correlations were generally stronger in the temporal cortex compared to the frontal cortex and stronger in the AD or in all subjects combined compared to the controls (**Table 2**). Some of the associations were lost upon adjusting for age- and sex in the multivariable analysis, especially the temporal cortex. However, the strong positive correlation of UFSP2 with soluble total tau (in all groups), and the association of soluble UFM1 with pS396/404-tau (in AD and the combined cohort) as well of insoluble UFM1 with pS396/404-tau (AD cohort only) remained significant in both regions. in addition, in the combined cohort soluble UFM1 was significantly associated with insoluble pS396/404-tau, while insoluble UFM1 was significantly associated with soluble total tau.

**Figure 3:**
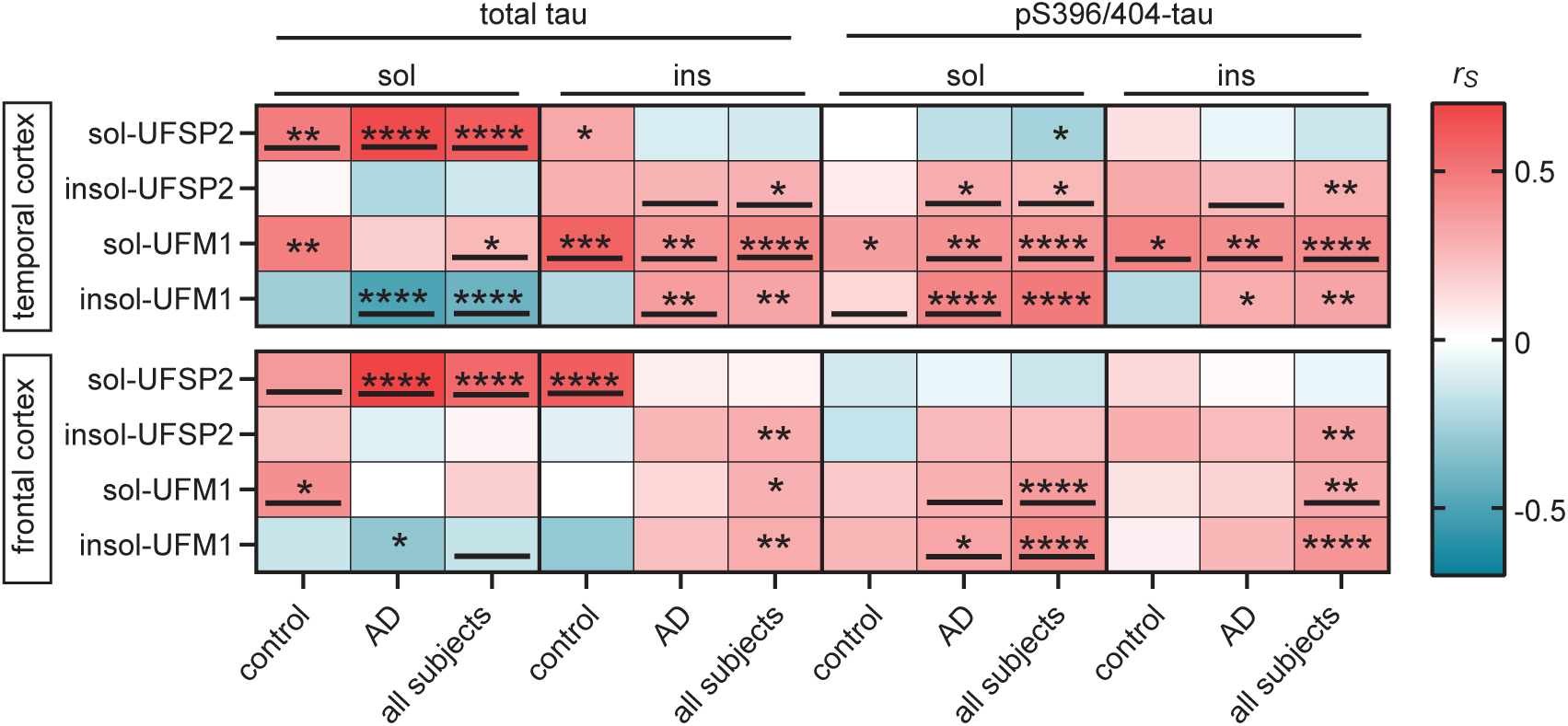
Soluble UFSP2 and insoluble UFM1 correlate with total and pS396/404-tau, respectively. (**A**) Heatmap of Spearman correlation coefficients (r_s_) illustrating significant correlations of UFM1 and UFSP2 protein level with total and pS396/404-tau levels in the temporal and frontal cortex of controls (n=41), AD (n=72) or combined groups (control + AD, n=113). A significance level of P<0.0125 after Bonferroni correction was used for the analysis: *P<0.0125, **P<0.0025, ***P < 0.00025, ****P<0.0001. Significant correlations that were confirmed by multivariable linear regression analysis (Table 2) have been underlined.

**Table 2:** Association of UFSP2 and UFM1 with tau.

| Cortex/Group/Variable | Association with sol UFSP2 |  | Association with ins UFSP2 |  | Association with sol UFM1 |  | Association with ins UFM1 |  |
| --- | --- | --- | --- | --- | --- | --- | --- | --- |
| | $\beta$ (95% CI) | P-value | $\beta$ (95% CI) | P-value | $\beta$ (95% CI) | P-value | $\beta$ (95% CI) | P-value |
| **Temporal Cortex** |  |  |  |  |  |  |  |  |
| Controls (N=41) |  |  |  |  |  |  |  |  |
| sol total tau | **1.03 (0.37, 1.68)** | **0.0030** | 2.14 (-7.26, 11.54) | 0.6463 | 19.19 (1.51, 36.87) | 0.0343 | -13.60 (-27.97, 0.76) | 0.0627 |
| ins total tau | 2.96 (0.28, 5.63) | 0.0315 | 30.11 (-4.77, 64.99) | 0.0884 | **105.86 (42.77, 168.95)** | **0.0017** | -52.55 (-107.97, 2.86) | 0.0623 |
| sol pS396/404-tau | -1.22 (-5.24, 2.81) | 0.5433 | -7.85 (-59.18, 43.47) | 0.7579 | 50.96 (-50.40, 152.33) | 0.3144 | **125.96 (55.88, 196.03)** | **0.0009** |
| ins pS396/404-tau | 0.99 (-1.20, 3.18) | 0.3644 | 18.33 (-9.12, 45.78) | 0.1839 | **74.00 (23.69, 124.31)** | **0.0051** | -27.07 (-71.22, 17.07) | 0.2214 |
| AD patients (N=72) |  |  |  |  |  |  |  |  |
| sol total tau | **1.67 (1.25, 2.09)** | **<0.0001** | -12.59 (-22.55, -2.64) | 0.0140 | 17.65 (-5.15, 40.45) | 0.1270 | **-15.56 (-24.10, -7.03)** | **0.0005** |
| ins total tau | -0.43 (-1.28, 0.42) | 0.3170 | **26.32 (12.76, 39.88)** | **0.0002** | **62.52 (32.84, 92.19)** | **<0.0001** | **16.82 (3.98, 29.66)** | **0.0111** |
| sol pS396/404-tau | -0.47 (-1.11, 0.17) | 0.1445 | **23.95 (14.17, 33.73)** | **<0.0001** | **58.61 (37.71, 79.51)** | **<0.0001** | **18.39 (9.19, 27.59)** | **0.0002** |
| ins pS396/404-tau | -0.33 (-1.31, 0.64) | 0.4978 | **22.54 (6.19, 38.90)** | **0.0076** | **64.09 (29.07, 99.10)** | **0.0005** | 11.70 (-3.53, 26.94) | 0.1298 |
| All subjects (N=113) |  |  |  |  |  |  |  |  |
| sol total tau | **1.39 (0.77, 2.02)** | **<0.0001** | -5.11 (-19.55, 9.33) | 0.4880 | **18.62 (5.05, 32.19)** | **0.0072** | **-15.04 (-22.21, -7.87)** | **<0.0001** |
| ins total tau | 1.03 (-2.25, 4.32) | 0.5376 | **26.84 (14.46, 39.21)** | **<0.0001** | **75.59 (36.60, 114.58)** | **0.0001** | -12.80 (-80.05, 54.46) | 0.7092 |
| sol pS396/404-tau | -0.49 (-1.11, 0.13) | 0.1197 | 17.72 (-7.03, 42.47) | 0.1605 | **58.29 (38.21, 78.36)** | **<0.0001** | 66.73 (-38.14, 171.60) | 0.2123 |
| ins pS396/404-tau | -0.02 (-1.12, 1.08) | 0.9718 | **21.41 (7.69, 35.14)** | **0.0022** | **67.39 (39.34, 95.45)** | **<0.0001** | -2.34 (-38.86, 34.19) | 0.9002 |
| **Frontal Cortex** |  |  |  |  |  |  |  |  |
| Controls (N=41) |  |  |  |  |  |  |  |  |
| sol total tau | **1.41 (0.34, 2.49)** | **0.0112** | 1.40 (-14.20, 16.99) | 0.8569 | **19.63 (5.41, 33.84)** | **0.0082** | -12.26 (-30.57, 6.05) | 0.1827 |
| ins total tau | **13.50 (6.85, 20.15)** | **0.0002** | -25.78 (-132.94, 81.37) | 0.6283 | -9.92 (-118.26, 98.42) | 0.8535 | -146.27 (-265.63, -26.91) | 0.0178 |
| sol pS396/404-tau | -2.17 (-5.65, 1.31) | 0.2140 | -21.07 (-67.72, 25.59) | 0.3656 | 16.32 (-30.95, 63.59) | 0.4879 | 32.84 (-22.89, 88.57) | 0.2397 |
| ins pS396/404-tau | 0.19 (-10.14, 10.52) | 0.9704 | 107.28 (-24.73, 239.30) | 0.1079 | -8.83 (-146.96, 129.30) | 0.8975 | -22.60 (-187.46, 142.26) | 0.7824 |
| AD patients (N=72) |  |  |  |  |  |  |  |  |
| sol total tau | **2.14 (1.57, 2.72)** | **<0.0001** | -5.15 (-15.31, 5.00) | 0.3142 | -5.53 (-20.81, 9.75) | 0.4723 | -11.92 (-22.19, -1.64) | 0.0237 |
| ins total tau | 1.02 (-0.13, 2.16) | 0.0809 | 14.94 (0.06, 29.82) | 0.0492 | 22.25 (-0.08, 44.57) | 0.0508 | 5.90 (-10.06, 21.86) | 0.4630 |
| sol pS396/404-tau | -0.06 (-1.05, 0.92) | 0.8968 | 12.49 (-0.04, 25.03) | 0.0508 | **46.44 (30.86, 62.01)** | **<0.0001** | **19.72 (7.14, 32.30)** | **0.0026** |
| ins pS396/404-tau | 0.71 (-0.55, 1.97) | 0.2637 | 16.67 (0.56, 32.78) | 0.0428 | **31.35 (7.65, 55.05)** | **0.0103** | 7.69 (-9.59, 24.96) | 0.3774 |
| All subjects (N=113) |  |  |  |  |  |  |  |  |
| sol total tau | **1.91 (1.25, 2.58)** | **<0.0001** | -3.16 (-11.47, 5.15) | 0.4559 | 7.24 (-17.41, 31.89) | 0.5650 | **-12.00 (-20.76, -3.24)** | **0.0072** |
| ins total tau | 6.84 (-5.36, 19.05) | 0.2719 | 14.14 (-0.32, 28.61) | 0.0553 | 20.89 (-0.55, 42.34) | 0.0561 | -59.04 (-206.55, 88.48) | 0.4328 |
| sol pS396/404-tau | -0.48 (-2.12, 1.16) | 0.5680 | 3.00 (-26.64, 32.63) | 0.8429 | **39.37 (14.35, 64.39)** | **0.0020** | **20.37 (8.34, 32.41)** | **0.0009** |
| ins pS396/404-tau | 0.70 (-0.52, 1.93) | 0.2612 | 39.00 (-37.53, 115.54) | 0.3179 | **30.17 (7.26, 53.08)** | **0.0099** | 7.35 (-9.51, 24.20) | 0.3929 |

sol=soluble; ins=insoluble; $\beta$ =regression coefficient; CI=confidence intervals. $\beta$ values, 95% CIs, and P-values for the separate control and AD groups result from multivariate regression models. $\beta$ values are interpreted as the increase in mean UFSP2 (on the square root scale when examining sol UFSP2) corresponding to a 1SD increase for the given continuous variables, which were examined on the untransformed, square root, or cube root scale. Models for controls were adjusted for age, sex, Braak stage, and Thal phase, and models for AD patients were adjusted for age, sex, presence of APOE ε4, Braak stage, and Thal phase. $\beta$ values, 95% CIs, and P-values for the analysis of all subjects results from a random effects meta-analysis combining the separate results from the control and AD groups. P-value < 0.0125 is considered as significant after applying a Bonferroni correction for multiple testing separately for each cortex and each disease group. Significant associations are shown in bold.

### UFSP2 KO enhances neuronal survival against DNA Damage through UFM1-dependent mechanism

In order to identify potential consequences of aberrant UFMylation, we first focused on the DNA damage response pathway that is known to be regulated by UFM1[13, 14, 37, 38]. AD neurons present with an abnormal accumulation of DNA lesions, suggesting that the DNA damage response is compromised in AD brains[22, 39, 40]. Utilizing available gene expression data from the temporal cortex of the same AD cases (n=72), we conducted a correlation analysis between the levels of UFM1 and UFSP2 proteins with the expression levels of DNA damage-related genes. Notably, soluble UFSP2 exhibited a significant correlation with 22 out of 37 genes (**Fig 4A, Table S10**). These association were spread across different sub-pathways and no single repair pathway stood out. This suggests a pivotal role of soluble UFSP2 for the DNA damage response within the human AD brain. In contrast, neither in-/soluble UFM1 nor insoluble UFSP2 showed a strong correlation with DNA damage response genes.

**Figure 4:**
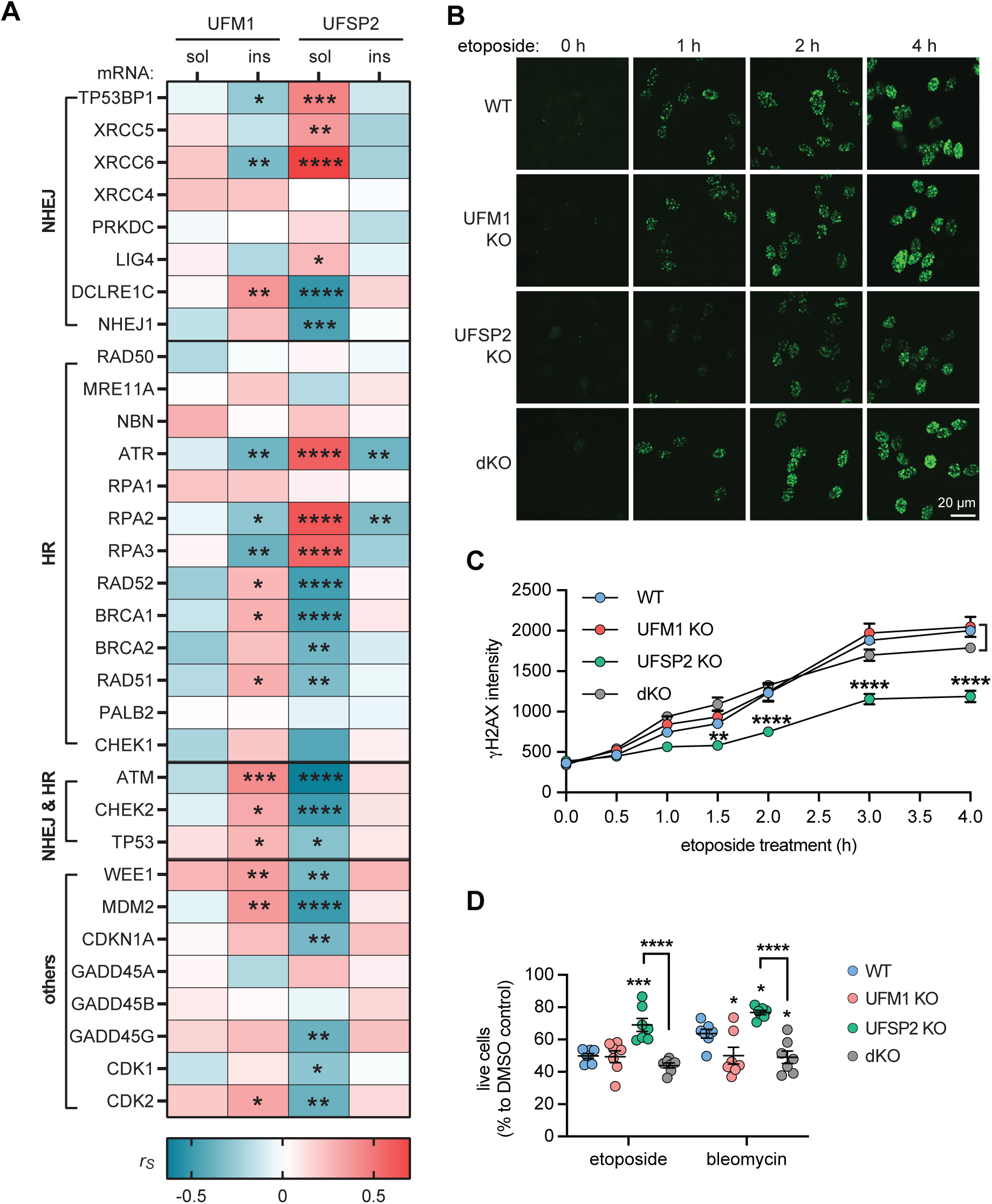
UFSP2 KO protects against DNA damage. (**A**) Heatmap of Spearman correlation coefficients (r_s_) illustrating correlations of soluble (sol) and insoluble (ins) UFM1 and UFSP2 protein levels, respectively, with the mRNA level of DNA damage related genes in the temporal cortex of AD subjects (n=72). mRNA levels were obtained by bulk RNAseq. A spearman’s test with significance level of P<0.05 was used for the analysis: *P<0.05, **P<0.01, ***P < 0.001, ****P<0.0001. See Supplementary Table S10 for r_s_ and p values. **(B, C)** Differentiated neurons with WT, UFM1 KO, UFSP KO or UFM1 and UFSP2 double KO (dKO) were treated with 10 µM etoposide for the indicated times and stained for γH2AX (green). (B) Representative microscope images at the indicated time points are shown for each genotype. Scale bars: 20 μm. (C) Images were analyzed by high content imaging for γH2AX intensity. Three independent experiments with multiple wells each were quantified over time. Data is shown as mean ± SEM. Statistical significance was assessed with two-way ANOVA. Shown is the least significant comparison for UFSP2 KO neurons when compared against any of the other three genotypes: **P<0.01, ****P<0.0001. (**D**) Percentage of live neurons (WT, UFM1 KO, UFSP2 KO, dKO) upon treatment with 100 µM etoposide and 10 µM bleomycin for 72 h. Cells were stained with a viability dyes and imaged. Live cells were identified and quantified by high content imaging. The live cell count of the stressed neurons was normalized to the live cell count of DMSO-treated cells for each genotype. Shown is the mean ± SEM of 7 independent experiments. Statistical significance to WT was assessed by one-way ANOVA followed by Dunnett’s test: *P<0.05, ***P<0.001. Statistical significance between UFSP2 KO and dKO cells was determined by student’s t test: ****P<0.001.

Neurons are particularly prone to the accumulation of DNA damage, a vulnerability that stems from their substantial energy demands, high levels of transcriptional activity, and longevity[41]. In order to mimic our findings of aberrant UFMylation from post-mortem brain, we used UFSP2 KO neurons, which, similar to AD brain, display low (absent) levels of UFSP2 and high levels of (conjugated) UFM1. As controls we utilized isogenic wild-type (WT) cells, as well as UFM1 KO cells for normal and absent of total UFM1, respectively. Further we generated a double knockout where we disabled both UFSP2 and UFM1 (dKO). To induce DNA double strand breaks in differentiated neurons, we utilized etoposide[42–44] and evaluated the dynamics of γH2Ax foci formation, a marker for DNA breaks[45]. Post etoposide treatment, UFSP2 KO neurons had substantially lower γH2Ax foci intensity compared to WT neurons (**Fig 4B**). This was not observed in dKO neurons, which displayed γH2Ax foci intensities similar to WT, indicating that UFSP2 KO neurons exhibit enhanced resistance to DNA damage in a UFM1-dependent manner. Interestingly, the formation of γH2Ax foci was not affected by loss of UFSP2 in undifferentiated neural progenitor cells (**Fig S5A**), suggesting that this might be a neuron-specific effect.

We next surveyed the viability of the neurons upon DNA damage and used etoposide or bleomycin to induce strand breaks. In line with the findings above, UFSP2 KO neurons showed greater survival upon DNA damage in comparison to WT neurons (**Fig 4B**). This advantage was negated by additionally knocking out UFM1 in UFSP2 KO neurons, indicating that the survival benefit of UFSP2 KO neurons is reliant on UFM1. Interestingly, UFM1 KO neurons showed no significant difference or even lower survival compared to WT neurons. Furthermore, the beneficial effect of UFPS2 KO seemed to be specific for differentiated neurons since neural progenitors did not show the same effects on survival (**Fig S5B**).

### UFSP2 Knockout modulates the unfolded protein response and neuronal survival under ER stress conditions

The UFMylation pathway also plays a central role in ER stress and its related unfolded protein response in mammals and plants[7, 11, 46]. Furthermore, the unfolded protein response is activated in AD and presents a target for therapy[21, 47–49]. To explore whether the aberrant UFMylation observed in AD could lead to an impaired unfolded protein response, we first examined the relationship between the levels of both soluble and insoluble UFSP2 and UFM1 proteins and unfolded protein response genes. Notably, expression of soluble UFSP2 was significantly associated with expression of five out of seven unfolded protein response genes. Of those, 4 (EIF2AK3/PERK, ATF4, DDIT3/CHOP and ERN1/IRE1α) were negatively correlated and one (ATF6) was positively correlated (**Fig 5A, Table S11**). This indicates that soluble UFSP2 might play an important role for the unfolded protein response in AD.

**Figure 5:**
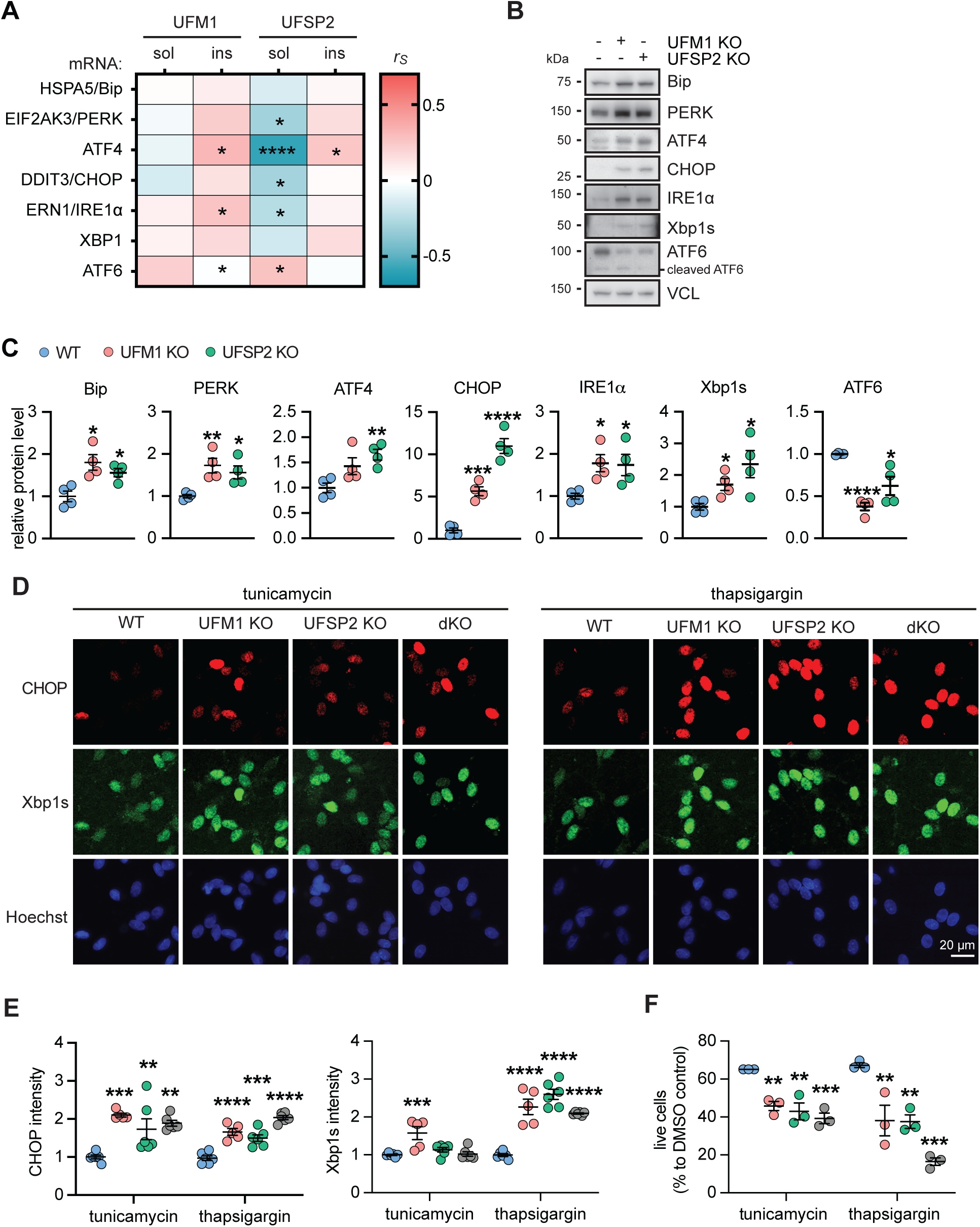
UFSP2 KO neurons exhibit a stronger unfolded protein response and higher susceptibility towards ER stress. (**A**) Heatmap of Spearman correlation coefficients (r_s_) illustrating significant correlations of soluble (sol) and insoluble (ins) UFM1 and UFSP2 protein levels, respectively, with the mRNA level of unfolded protein response related genes in the temporal cortex of AD subjects (n=72). mRNA levels were obtained by bulk RNAseq. A significance level of P<0.05 was used for the analysis: *P<0.05, **P<0.01, ***P< 0.001, ****P<0.0001. See Supplementary Table S11 for r_s_ and p values. **(B, C)** Immunoblot analysis and quantification of expression of seven unfolded protein response related proteins in differentiated neurons with UFM1 KO, UFSP2 KO or a double KO (dKO) compared to isogenic controls (WT). Shown is the normalized mean ± SEM from four independent experiments. **(D, E)** Neurons (WT, UFM1 KO, UFSP2 KO or UFM1/UFSP2 double KO (dKO)) were treated with tunicamycin or thapsigargin for 16 hours, and then fixed and stained with anti-CHOP (red) and anti-Xbp1s (green) antibodies. Nuclei were labeled with Hoechst 33342 (blue). (D) Representative images are shown for untreated or treated neurons for each genotypes. Scale bars: 20 μm. (E) Images were analyzed by high content imaging for CHOP and Xbp1s intensity, respectively. Data is shown as mean ± SEM from n=5-6 independent experiments. Statistical comparison to WT was assessed with one-way ANOVA followed by Dunnett’s posthoc test: **P<0.01, ***P<0.001, ****P<0.0001. **(F)** Percentage of live neurons (WT, UFM1 KO, UFSP2 KO or UFM1/UFSP2 double KO (dKO)) upon treatment with tunicamycin of thapsigargin for 72 h. Cells were fixed and stained with viability/cytotoxicity dyes, imaged and analyzed by high content imaging. The number of live cells was normalized to the cell count of DMSO-treated cells for each genotype. Shown is the mean ± SEM of 3 independent experiments. Statistical significance was assessed with one-way ANOVA followed by Dunnett’s post-hoc test: **P<0.01, ***P<0.001.

Next, we aimed to explore the impact of UFSP2 KO on the unfolded protein response pathway in neurons and analyzed the protein levels of seven unfolded protein response molecules in WT, UFM1 KO, and UFSP2 KO neurons at baseline, in the absence of stress (**Fig 5B**). This revealed remarkable differences between WT and UFSP2 KO neurons for each of the investigated proteins. Consistent with the mostly negative correlation between UFSP2 protein and gene expression levels in AD brain, Bip, PERK, ATF4, CHOP, IRE1α, and Xbp1s were all increased in UFSP2 KO neurons, while ATF6 levels were decreased (**Fig 5B,C**). Notably, only the full-length ATF6 protein, not the cleaved ATF6 which is the active form upon ER stress[50], showed a decrease, suggesting that this reduction is not a result of heightened unfolded protein response activation.

We next induced ER stress with tunicamycin or thapsigargin[51] and monitored induction of Xbp1s and CHOP by high content imaging of cells that were stained by immunofluorescence. Following both treatments, levels of CHOP immunoreactivity were significantly higher in neurons with UFM1 KO, UFSP2 KO, or with dKO compared to WT cells (**Fig 5D, E**). The expression levels of Xbp1s were also significantly elevated in the same genotypes compared to WT, at least in response to thapsigargin. In order to assess the resilience of UFSP2 and UFM1 KO neurons against ER stress we measured the survival. As expected, neurons with UFSP2 KO exhibited a significantly lower survival rate compared to WT after both treatments (**Fig 5F**). UFM1 KO also caused lower survival compared to WT but could not reverse the effect of the UFSP2 KO. This suggest that resilience towards ER stress is highly susceptible to increases and decreases of the UFMylation pathway and this might also affect the viability of neurons.

## DISCUSSION

The UFMylation pathway is implicated in a variety of biological processes known to be disrupted in AD, and deficiencies in this pathway have been linked to neurodevelopmental disorders[28–30, 52]. In addition, the UFM1 pathway was very recently identified as potent modulator of tau aggregation upon seeding[26, 53]. Therefore, the UFM1 pathway is of high relevance for AD. Yet, the specific role of UFMylation in the development and progression of AD remains elusive. Here, we comprehensively explored changes of the UFM1 pathway in AD. We utilized RNAseq data and performed a thorough biochemical analysis of UFM1 in two different post-mortem brain cohorts and across early and later affected brain regions. We correlated our findings with additional biochemical and genetic data and further validated findings in neurons upon genetic and pharmacological manipulation.

To explore changes in the UFM1 pathway, we first examined published single nuclei transcriptomic data[35] and discovered that most genes related to the UFMylation pathway were dysregulated in excitatory neurons of AD patients. Previous studies have reported that excitatory neurons are more susceptible to neurodegeneration[54]. Biochemical assessment of the UFMylation pathway in post-mortem brains revealed a solubility shift of UFSP2, while UFM1 levels were significantly elevated in both cortical areas in AD patients compared to controls. Importantly, consistent with the role as the UFM1 protease, neurons with UFSP2 KO showed a marked increase of conjugated UFM1. This finding not only reflects the negative correlation between UFM1 and UFSP2 observed in the AD brain but also suggests that UFSP2 KO neurons could serve as a relevant model to study the aberrant UFMylation observed in AD.

Our results showed that total UFM1 was abnormally accumulated in AD brain. In the absence of alterations in free UFM1, this change represents hyperUFMylation, an increase in specifically conjugated UFM1. It is possible that there is a general increase of UFM1 attached as monomer or in chains on one or several physiological substrates. Most knowledge about UFM1 substrates is derived from studies in cancer cell lines[15, 27, 55, 56]. Alternatively, in AD UFM1 could accumulate on a substrate that is normally not modified by UFM1 and further studies are needed to investigate targets of physiological and pathological UFMylation in the brain. We identified a strong correlation of UFM1 levels with pathological tau, suggesting that hyperUFMylation might be linked to tau pathogenesis. This is in line with recent studies that identified suppression of the UFMylation cascade as potent inhibitor of tau aggregation and seeding in human induced pluripotent stem cell (iPSC)-derived neurons and tau transgenic mice[26]. However, the mechanism of this interaction remains elusive. It is not known whether UFM1 modifies tau or other substrates that may affect tau aggregation through pathways such as aberrant ER stress, ER-phagy, and ribosomal quality control.

To explore whether abnormalities of UFMylation are influenced by AD disease progression, we specifically examined the expression of UFM1 and UFSP2 and their correlation with tau in the earlier-affected superior temporal cortex and the later-affected frontal cortex. Both brain regions exhibited higher levels of UFM1 and insoluble UFSP2 in AD. A notable difference was that soluble UFSP2 was significantly reduced in the temporal cortex, whereas in the frontal cortex there was only a trend. It is therefore unclear whether the loss of the deUFMylation enzyme UFSP2 causes the hyperUFMylation or if hyperUFMylation is induced by the presence of a substrate, such as tau and the loss of soluble UFSP2 might further contribute to it. Moreover, UFM1 was positively correlated with several pathological forms of tau in the temporal cortex, whereas in the frontal cortex, it only correlated positively with soluble pS396/404-tau. It is conceivable, that this is caused by incomplete pathological changes in tau in this later-affected region, while changes in UFM1 are already observed. This would place UFM1 parallel or upstream of pathogenic tau changes. On the other hand, UFM1 could also be affected by tau deposition itself, in line with a shift towards insoluble UFM1 in tau seeded iPSC-derived neurons[26]. Nevertheless, insoluble UFM1 was negatively correlated with soluble UFSP2 in both AD and in controls in both brain regions, suggesting that this relationship may be universal and unaffected by AD presence and progression and that increasing UFSP2 activity might be a good strategy to combat hyperUFMylation.

In order to further determine the functional effects of hyperUFMylation, we created UFSP2 KO cells and tested functional effects of DNA damage and unfolded protein response in neurons. In the context of DNA damage, we found that levels of soluble UFSP2 correlated with the expression of a majority of DNA damage response-related genes. Surprisingly, UFSP2 KO neurons displayed a reduced sensitivity to DNA damage, exhibiting a milder DNA damage response compared to wild-type, a phenomenon reliant on the accumulation of UFM1-modified proteins. This suggests hyperUFMylation might confer a protective effect against DNA damage in AD neuronal cells. This is in contrast to cancer cells, where UFSP2 KO is known to enhance DNA damage response to counteract DNA damage[38]. This discrepancy could be attributed to the fundamental differences between non-proliferating neurons and proliferating cancer cells, further highlighting the need to study the UFM1 pathway in neurons and in the brain.

In the context of the unfolded protein response, we found a significant correlation between levels of soluble UFSP2 and the mRNA expression of numerous unfolded protein response genes in human temporal cortex of AD brain. Consistently, without any treatment, the expression levels of six key unfolded protein response proteins were elevated in UFSP2 KO neurons, indicating an inherently higher unfolded protein response in UFSP2 KO neurons compared to WT at baseline. Moreover, our findings reveal that UFSP2 KO neurons exhibit increased sensitivity to ER stress as they showed higher levels of CHOP following tunicamycin or thapsigargin treatment, implicating a pronounced unfolded protein response activation. Given that ER stress-induced apoptosis is predominantly mediated by CHOP[21, 57], this could account for the observed reduction in survival rates. Our results therefore indicate that a reduction in soluble UFSP2 levels may be a key factor in the continuous activation of the unfolded protein response in AD brain[21]. However, the susceptibility towards ER stress was not only increased by UFSP2 but also by UFM1 KO, highlighting that hyper-as well as hypoUFMylation both can have negative effects on the survival of neurons in certain contexts.

The main limitation of this study is the relatively small sample size, which results in a lack of power to detect differences and associations. In particular, the control group is not very large. Therefore, the possibility of a type II error (i.e., a false-negative finding) is important to consider, and we cannot conclude that a true difference does not exist simply due to the occurrence of a non-significant p-value in our study.

Collectively, our data indicates that increasing UFSP2 activity might be an attractive target to counteract the observed hyperUFMylation that is linked to pathological tau in AD brain. However, it should be noted that the loss of UFMylation might increase the unfolded protein response and might have further far-reaching effects. The loss of UFM1 has been connected to severe neurodevelopmental phenotypes[15, 27, 31] and therefore unintended consequences of such approach will have to be carefully monitored. Our study underscores the critical need to identify specific substrates and molecular mechanisms of UFM1 in cell culture and animal models, to identify risks and benefits of manipulating the UFM1 pathway as potential therapeutic avenue for AD.

## Abbreviations

Aβ: beta-amyloid
MMSE: mini mental state examination
M²OVE-AD: Molecular Mechanisms of the Vascular Etiology of AD
TBS: Tris-buffered saline
TX: 1% Triton X-100 in TBS
FA: formic acid
RIPA: radioimmunoprecipitation assay
RT: room temperature
EGF: epidermal growth factor
FGF: fibroblasts growth factor
KO: knock out
dKO: double knockouts
PCR: polymerase chain reaction
MSD: Meso Scale Discovery
ELISA: enzyme-linked immunosorbent assay
ER: Endoplasmic reticulum
IPSC: induced pluripotent stem cell

## Declarations

### Ethics approval

All brain samples are from autopsies performed after approval by the legal next-of-kin. Research on de-identified postmortem brain tissue is considered exempt from human subjects regulations by the Mayo Clinic Institutional Review Board.

### Consent for publication

Not applicable

### Availability of data and materials

The data that support the findings of this study are available from the corresponding author, upon reasonable request.

### Competing interests

The authors declare that they have no competing interests.

### Fundings

This work was supported by a Florida Department of Health Ed and Ethel Moore Alzheimer’s disease research program grant 22A07 (to F.C.F.), NIH grants R56 AG062556 (to W.S.), and a fellowship from Alzheimer’s Association AARF-22-974258 (to T.Y.). This work was supported by the Mayo Clinic Office of Research Equity, Inclusion and Diversity and Center for Biomedical Discovery.

### Authors’ contributions

T.Y., W. S. and F.C.F. conceived and designed the study; T.Y. and F.C.F. performed the experiments and analyzed the data; M.G.H. and E.C. analyzed the data; M.E.M. and D.W.D provided the post-mortem tissues; B.D.R. cut the post-mortem tissues; C.L., and G.B., provided the protein expression data from M^2^OVE-AD cohort; X.W. and N.T. provided bulk RNA-seq data from M^2^OVE-AD cohort; Z.L. participated the design of the DNA damage related experiments; T.Y., W.S. and F.C.F. wrote the manuscript; All authors discussed the results and commented on the manuscript. All authors read and approved the final manuscript.

## Acknowledgments

We thank all patients and family members who made this study possible. We are grateful to the late Dr. Peter Davies for providing the pS396/404-tau (PHF1) antibodies.

## Notes

### Competing Interest Statement

The authors have declared no competing interest.

## REFERENCES

1. Komatsu M, Chiba T, Tatsumi K, Iemura S, Tanida I, Okazaki N, Ueno T, Kominami E, Natsume T, Tanaka K: A novel protein-conjugating system for Ufm1, a ubiquitin-fold modifier. Embo j 2004, 23:1977–1986.

2. Sasakawa H, Sakata E, Yamaguchi Y, Komatsu M, Tatsumi K, Kominami E, Tanaka K, Kato K: Solution structure and dynamics of Ufm1, a ubiquitin-fold modifier 1. Biochemical and Biophysical Research Communications 2006, 343:21–26.

3. Kang SH, Kim GR, Seong M, Baek SH, Seol JH, Bang OS, Ovaa H, Tatsumi K, Komatsu M, Tanaka K, Chung CH: Two novel ubiquitin-fold modifier 1 (Ufm1)-specific proteases, UfSP1 and UfSP2. J Biol Chem 2007, 282:5256–5262.

4. Mizushima T, Tatsumi K, Ozaki Y, Kawakami T, Suzuki A, Ogasahara K, Komatsu M, Kominami E, Tanaka K, Yamane T: Crystal structure of Ufc1, the Ufm1-conjugating enzyme. Biochemical and Biophysical Research Communications 2007, 362:1079–1084.

5. Bacik J-P, Walker JR, Ali M, Schimmer AD, Dhe-Paganon S: Crystal structure of the human ubiquitin-activating Enzyme 5 (UBA5) bound to ATP: MECHANISTIC INSIGHTS INTO A MINIMALISTIC E1 ENZYME. J Biol Chem 2010, 285:20273–20280.

6. Tatsumi K, Sou Y-s, Tada N, Nakamura E, Iemura S-i, Natsume T, Kang SH, Chung CH, Kasahara M, Kominami E, et al: A novel type of E3 ligase for the Ufm1 conjugation system. J Biol Chem 2010, 285:5417–5427.

7. Lemaire K, Moura RF, Granvik M, Igoillo-Esteve M, Hohmeier HE, Hendrickx N, Newgard CB, Waelkens E, Cnop M, Schuit F: Ubiquitin fold modifier 1 (UFM1) and its target UFBP1 protect pancreatic beta cells from ER stress-induced apoptosis. PLOS ONE 2011, 6:e18517.

8. Peter JJ, Magnussen HM, DaRosa PA, Millrine D, Matthews SP, Lamoliatte F, Sundaramoorthy R, Kopito RR, Kulathu Y: A non-canonical scaffold-type E3 ligase complex mediates protein UFMylation. The EMBO Journal 2022, 41:e111015.

9. Liang Q, Jin Y, Xu S, Zhou J, Mao J, Ma X, Wang M, Cong Y-S: Human UFSP1 translated from an upstream near-cognate initiation codon functions as an active UFM1-specific protease. J Biol Chem 2022:102016.

10. Millrine D, Cummings T, Matthews SP, Peter JJ, Magnussen HM, Lange SM, Macartney T, Lamoliatte F, Knebel A, Kulathu Y: Human UFSP1 is an active protease that regulates UFM1 maturation and UFMylation. Cell Reports 2022, 40:111168.

11. Zhang Y, Zhang M, Wu J, Lei G, Li H: Transcriptional regulation of the Ufm1 conjugation system in response to disturbance of the endoplasmic reticulum homeostasis and inhibition of vesicle trafficking. PLOS ONE 2012, 7:e48587.

12. Liu J, Wang Y, Song L, Zeng L, Yi W, Liu T, Chen H, Wang M, Ju Z, Cong Y-S: A critical role of DDRGK1 in endoplasmic reticulum homoeostasis via regulation of IRE1α stability. Nature Communications 2017, 8:14186.

13. Qin B, Yu J, Nowsheen S, Wang M, Tu X, Liu T, Li H, Wang L, Lou Z: UFL1 promotes histone H4 ufmylation and ATM activation. Nature Communications 2019, 10:1242.

14. Wang Z, Gong Y, Peng B, Shi R, Fan D, Zhao H, Zhu M, Zhang H, Lou Z, Zhou J, et al: MRE11 UFMylation promotes ATM activation. Nucleic Acids Research 2019, 47:4124–4135.

15. Zhou X, Mahdizadeh SJ, Le Gallo M, Eriksson LA, Chevet E, Lafont E: UFMylation: a ubiquitin-like modification. Trends in Biochemical Sciences 2023.

16. Gerakis Y, Quintero M, Li H, Hetz C: The UFMylation system in proteostasis and beyond. Trends in Cell Biology 2019, 29:974–986.

17. DeJesus R, Moretti F, McAllister G, Wang Z, Bergman P, Liu S, Frias E, Alford J, Reece-Hoyes JS, Lindeman A, et al: Functional CRISPR screening identifies the ufmylation pathway as a regulator of SQSTM1/p62. eLife 2016, 5:e17290.

18. Liang JR, Lingeman E, Luong T, Ahmed S, Muhar M, Nguyen T, Olzmann JA, Corn JE: A genome-wide ER-phagy screen highlights key roles of mitochondrial metabolism and ER-resident UFMylation. Cell 2020, 180:1160–1177.e1120.

19. Xi P, Ding D, Zhou J, Wang M, Cong Y-S: DDRGK1 regulates NF-κB activity by modulating IκBα stability. PLOS ONE 2013, 8:e64231.

20. Tao Y, Yin S, Liu Y, Li C, Chen Y, Han D, Huang J, Xu S, Zou Z, Yu Y: UFL1 promotes antiviral immune response by maintaining STING stability independent of UFMylation. Cell Death & Differentiation 2022.

21. Ajoolabady A, Lindholm D, Ren J, Pratico D: ER stress and UPR in Alzheimer’s disease: mechanisms, pathogenesis, treatments. Cell Death & Disease 2022, 13:706.

22. Dileep V, Boix CA, Mathys H, Marco A, Welch GM, Meharena HS, Loon A, Jeloka R, Peng Z, Bennett DA, et al: Neuronal DNA double-strand breaks lead to genome structural variations and 3D genome disruption in neurodegeneration. Cell 2023, 186:4404–4421.e4420.

23. Di Meco A, Curtis ME, Lauretti E, Praticò D: Autophagy dysfunction in Alzheimer’s disease: mechanistic insights and new therapeutic opportunities. Biological Psychiatry 2020, 87:797–807.

24. Leng F, Edison P: Neuroinflammation and microglial activation in Alzheimer disease: where do we go from here? Nature Reviews Neurology 2021, 17:157–172.

25. Wilson DM, Cookson MR, Van Den Bosch L, Zetterberg H, Holtzman DM, Dewachter I: Hallmarks of neurodegenerative diseases. Cell 2023, 186:693–714.

26. Parra Bravo C, Giani AM, Perez JM, Zhao Z, Wan Y, Samelson AJ, Wong MY, Evangelisti A, Cordes E, Fan L, et al: Human iPSC 4R tauopathy model uncovers modifiers of tau propagation. Cell 2024.

27. Wang X, Xu X, Wang Z: The post-translational role of UFMylation in physiology and disease. Cells 2023, 12:2543.

28. Nahorski MS, Maddirevula S, Ishimura R, Alsahli S, Brady AF, Begemann A, Mizushima T, Guzmán-Vega FJ, Obata M, Ichimura Y, et al: Biallelic UFM1 and UFC1 mutations expand the essential role of ufmylation in brain development. Brain 2018, 141:1934–1945.

29. Colin E, Daniel J, Ziegler A, Wakim J, Scrivo A, Haack Tobias B, Khiati S, Denommé A-S, Amati-Bonneau P, Charif M, et al: Biallelic variants in UBA5 reveal that disruption of the UFM1 cascade can result in early-onset encephalopathy. The American Journal of Human Genetics 2016, 99:695–703.

30. Ni M, Afroze B, Xing C, Pan C, Shao Y, Cai L, Cantarel BL, Pei J, Grishin NV, Hewson S, et al: A pathogenic UFSP2 variant in an autosomal recessive form of pediatric neurodevelopmental anomalies and epilepsy. Genetics in Medicine 2021.

31. Yang S, Moy N, Yang R: The UFM1 conjugation system in mammalian development. Developmental Dynamics 2023, 252:976–985.

32. Liu C-C, Yamazaki Y, Heckman MG, Martens YA, Jia L, Yamazaki A, Diehl NN, Zhao J, Zhao N, DeTure M, et al: Tau and apolipoprotein E modulate cerebrovascular tight junction integrity independent of cerebral amyloid angiopathy in Alzheimer’s disease. Alzheimer’s & Dementia 2020, 16:1372–1383.

33. Fiesel FC, Fričová D, Hayes CS, Coban MA, Hudec R, Bredenberg JM, Broadway BJ, Markham BN, Yan T, Boneski PK, et al: Substitution of PINK1 Gly411 modulates substrate receptivity and turnover. Autophagy 2022:1–22.

34. DerSimonian R, Laird N: Meta-analysis in clinical trials. Controlled Clinical Trials 1986, 7:177–188.

35. Mathys H, Davila-Velderrain J, Peng Z, Gao F, Mohammadi S, Young JZ, Menon M, He L, Abdurrob F, Jiang X, et al: Single-cell transcriptomic analysis of Alzheimer’s disease. Nature 2019.

36. Augustinack JC, Schneider A, Mandelkow E-M, Hyman BT: Specific tau phosphorylation sites correlate with severity of neuronal cytopathology in Alzheimer’s disease. Acta Neuropathol 2002, 103:26–35.

37. Liu J, Guan D, Dong M, Yang J, Wei H, Liang Q, Song L, Xu L, Bai J, Liu C, et al: UFMylation maintains tumour suppressor p53 stability by antagonizing its ubiquitination. Nature Cell Biology 2020, 22:1056–1063.

38. Qin B, Yu J, Zhao F, Huang J, Zhou Q, Lou Z: Dynamic recruitment of UFM1-specific peptidase 2 to the DNA double-strand breaks regulated by WIP1. Genome Instability & Disease 2022, 3:217–226.

39. Silva ART, Santos ACF, Farfel JM, Grinberg LT, Ferretti REL, Campos AHJFM, Cunha IW, Begnami MD, Rocha RM, Carraro DM, et al: Repair of oxidative DNA damage, cell-cycle regulation and neuronal death may influence the clinical manifestation of Alzheimer’s disease. PLOS ONE 2014, 9:e99897.

40. Chen J, Cohen ML, Lerner AJ, Yang Y, Herrup K: DNA damage and cell cycle events implicate cerebellar dentate nucleus neurons as targets of Alzheimer’s disease. Mol Neurodegener 2010, 5:60.

41. Welch G, Tsai L-H: Mechanisms of DNA damage-mediated neurotoxicity in neurodegenerative disease. EMBO reports 2022, n/a:e54217.

42. Madabhushi R, Gao F, Pfenning Andreas R, Pan L, Yamakawa S, Seo J, Rueda R, Phan TX, Yamakawa H, Pao P-C, et al: Activity-induced DNA breaks govern the expression of neuronal early-response genes. Cell 2015, 161:1592–1605.

43. Mitra J, Guerrero EN, Hegde PM, Liachko NF, Wang H, Vasquez V, Gao J, Pandey A, Taylor JP, Kraemer BC, et al: Motor neuron disease-associated loss of nuclear TDP-43 is linked to DNA double-strand break repair defects. Proceedings of the National Academy of Sciences 2019, 116:4696–4705.

44. Asada-Utsugi M, Uemura K, Ayaki T, T. Uemura M, Minamiyama S, Hikiami R, Morimura T, Shodai A, Ueki T, Takahashi R, et al: Failure of DNA double-strand break repair by tau mediates Alzheimer’s disease pathology in vitro. Communications Biology 2022, 5:358.

45. Kinner A, Wu W, Staudt C, Iliakis G: γ-H2AX in recognition and signaling of DNA double-strand breaks in the context of chromatin. Nucleic Acids Research 2008, 36:5678–5694.

46. Li B, Niu F, Zeng Y, Tse MK, Deng C, Hong L, Gao S, Lo SW, Cao W, Huang S, et al: Ufmylation reconciles salt stress-induced unfolded protein responses via ER-phagy in Arabidopsis. Proceedings of the National Academy of Sciences 2023, 120:e2208351120.

47. Hoozemans JJM, Veerhuis R, Van Haastert ES, Rozemuller JM, Baas F, Eikelenboom P, Scheper W: The unfolded protein response is activated in Alzheimer’s disease. Acta Neuropathol 2005, 110:165–172.

48. Hetz C, Mollereau B: Disturbance of endoplasmic reticulum proteostasis in neurodegenerative diseases. Nature Reviews Neuroscience 2014, 15:233–249.

49. Yoon Sung O, Park Dong J, Ryu Jae C, Ozer Hatice G, Tep C, Shin Yong J, Lim Tae H, Pastorino L, Kunwar Ajaya J, Walton James C, et al: JNK3 perpetuates metabolic stress induced by Aβ peptides. Neuron 2012, 75:824–837.

50. Ye J, Rawson RB, Komuro R, Chen X, Davé UP, Prywes R, Brown MS, Goldstein JL: ER stress induces cleavage of membrane-bound ATF6 by the same proteases that process SREBPs. Molecular Cell 2000, 6:1355–1364.

51. Zohreh G, Patrick S, Benjamin F, Steven MC, David SP, Sean PC: Neuronal apoptosis induced by endoplasmic reticulum stress is regulated by ATF4–CHOP-mediated induction of the Bcl-2 homology 3-only member PUMA. The Journal of Neuroscience 2010, 30:16938.

52. Muona M, Ishimura R, Laari A, Ichimura Y, Linnankivi T, Keski-Filppula R, Herva R, Rantala H, Paetau A, Pöyhönen M, et al: Biallelic variants in UBA5 link dysfunctional UFM1 ubiquitin-like modifier pathway to severe infantile-onset encephalopathy. The American Journal of Human Genetics 2016, 99:683–694.

53. Samelson AJ, Ariqat N, McKetney J, Rohanitazangi G, Bravo CP, Goodness D, Tian R, Grosjean P, Abskharon R, Eisenberg D, et al: CRISPR screens in iPSC-derived neurons reveal principles of tau proteostasis. bioRxiv 2023:2023.2006.2016.545386.

54. Fu H, Possenti A, Freer R, Nakano Y, Hernandez Villegas NC, Tang M, Cauhy PVM, Lassus BA, Chen S, Fowler SL, et al: A tau homeostasis signature is linked with the cellular and regional vulnerability of excitatory neurons to tau pathology. Nature Neuroscience 2019, 22:47–56.

55. Chung CH, Yoo HM: Emerging role of protein modification by UFM1 in cancer. Biochemical and Biophysical Research Communications 2022, 633:61–63.

56. Jing Y, Mao Z, Chen F: UFMylation system: An emerging player in tumorigenesis. Cancers 2022, 14:3501.

57. Uddin MS, Yu WS, Lim LW: Exploring ER stress response in cellular aging and neuroinflammation in Alzheimer’s disease. Ageing Research Reviews 2021, 70:101417.

