## Supplemental figure and tables for "The UFMylation pathway is impaired in Alzheimer’s disease": UFMylation in AD_SuppInfo-20240524.pdf

Affiliations:

### SUPPLEMENTARY FIGURE LEGENDS

**Supplementary Figure 1: UFMylation gene expression is dysregulated in AD excitatory neurons**

Meta-analysis of published gene expression data derived from single nuclei transcriptomics from AD patients versus controls [35]. (**A**) Heatmap of relative gene expression changes of all eight UFMylation pathway components in excitatory neurons (EX), inhibitory neurons (IN), astrocytes (AS), oligodendrocytes (OL), oligodendrocyte precursor cells (OPC), and microglia (MG). (**B**) Heatmap of relative gene expression changes of ubiquitin-like (UBL) pathways components in excitatory neurons. Significance levels used are as follows: *: a gene meets 2-sided Wilcoxon-rank-sum test, FDR<0.01, |log-Fold-Change|>0.25, **: a gene meets 2-sided Wilcoxon-rank-sum test, FDR<0.01, |log-Fold-Change|>0.25, and Poisson mixed-model FDR<0.05. See Supplementary Table S4 for related data.

**Supplementary Figure 2: Optimization and assessments of the MSD ELISA for UFM1**

(**A**) Western blot confirming the absence of UFM1 in UFM1 KO neurons, which were generated by CRISPR-Cas9. Black arrowhead indicates monomeric UFM1, gray arrowheads point to conjugated UFM1 species. Asterisks denote non-specific bands. (**B**) Comparison of five commercial UFM1 antibodies using WT, UFM1 KO and UFSP2 KO neuron samples for western blot, respectively. Ab1 (Abcam, ab109305) showed a clear difference between WT and UFM1 cells as well as between WT and UFSP2 cells. Ab2 (Sigma, HPA039758), Ab3 (Protein Tech Group, 15883-1-AP) and Ab4 (LS Bio, LS-C807041) show additional bands in UFSP2 KO cells, suggesting that they can detect the abundant conjugated form of UFM1 in these cells but do not show a clear difference between WT and UFM1 KO cells. Ab5 (LS Bio, LS-C500000) seems unspecific since there is no difference between WT, UFM1 KO and UFSP2 KO cells. (**C**) MSD ELISA test with the same five commercial UFM1 antibodies using denatured lysates from WT, UFM1 KO and UFSP2 KO neurons. Only Ab1 is suitable for MSD ELISA. Statistical significance was assessed with one-way ANOVA. Asterisks indicate statistical difference to WT (****P< 0.0001). (**D**) Lysates from UFM1 KO neurons were mixed with lysates from UFSP2 KO neuron to generate lysates with different concentrations of UFM1. Shown is a linear regression of UFM1 MSD ELISA values against this range of UFM1 concentrations.

**Supplementary Figure 3: UFSP2 MSD ELISA**

(**A**) Western blot of wild-type (-) and three different neuronal clones in which UFSP2 had been knocked out using CRISPR-Cas9. All UFSP2 KO neurons show lower level of unconjugated (black arrowhead) but higher levels of conjugated UFM1 (gray arrowheads). (**B**) MSD ELISA to measure UFSP2 using denatured lysates from WT, UFM1, or UFSP2 KO neurons. (**C**) Spearman correlation of UFSP2 protein level in samples of the exploratory cohort (controls (blue) and AD (maroon)) measured by western blot and MSD ELISA. The correlation coefficient and P value are indicated. (**D**) Analysis of MSD ELISA-assessed soluble UFSP2 protein levels by disease group. Statistical analysis was performed by Mann-Whitney test: **P<0.01. (**E**) Spearman correlation of MSD ELISA-assessed soluble UFSP2 protein levels with soluble and insoluble total UFM1 levels indicate a strong, significant negative correlation for both. The correlation coefficient and P value are indicated.

**Supplementary Figure 4: Correlations of UFM1 and UFSP2 proteins with AD-related proteins that were biochemically measured from the temporal cortex**

Heatmap of Spearman correlation coefficients ($r_s$) illustrating the absence of significant correlations of soluble and insoluble UFM1 and UFSP2 with most AD markers except for tau. Levels of ApoE, Aβ40, Aβ42, tau, and pT231-tau were previously measured in a separate study from the same individuals using three fractions: tris buffered saline (TBS), Triton X-100 (TX) and formic acid (FA) [32]. A significant correlation of between soluble UFM1 and soluble total and pT231-tau in the TBS fraction, as well as insoluble UFM1 and pT231-tau in the TBS and TX fraction prompted us to investigate tau further. *P<0.05, **P<0.01.

**Supplementary Figure 5: Knock out of UFSP2 does not affect the DNA damage response in undifferentiated neuronal precursor cells.**

(**A**) Undifferentiated WT, UFM1 KO and UFSP2 KO neuronal precursors were treated with 10 µM etoposide for the indicated times and stained with an antibody against γH2AX. γH2AX intensity was measured by high content imaging. Intensity values were averaged and are presented as mean ± SEM from three independent experiments. Statistical significance was assessed with two-way ANOVA. No significant difference was identified. (**B**) Percentage of live neuronal precursor cells (WT, UFM1 KO, or UFSP2 KO) upon treatment with etoposide or bleomycin for 72 h. Cells were fixed and stained with viability/cytotoxicity dyes, imaged and analyzed by high content imaging. The number of live cells was normalized to the cell count of DMSO-treated cells for each genotype. Shown is the mean ± SEM of four independent experiments. Statistical significance was assessed with one-way ANOVA followed by Dunnett's post-hoc test: no significant difference was identified.

### LIST OF SUPPLEMENTARY TABLES

**Supplementary Table 1:** Subject characteristics of exploratory cohort.

**Supplementary Table 2:** Subject characteristics of main cohort

**Supplementary Table 3:** Transformation of variables for linear regression analyses

**Supplementary Table 4:** Differential single nuclei expression of UBL pathway components in samples with and without AD pathology

**Supplementary Table 5:** Correlation of UFM1 values with other UFMylation proteins in exploratory cohort

**Supplementary Table 6:** Correlation of UFSP2 with UFM1 in human brain

**Supplementary Table 7:** Associations of primary clinical and disease parameters with UFSP2 and UFM1

**Supplementary Table 8:** Correlations of UFSP2 and UFM1 with common AD-related parameters in temporal cortex of AD patients

**Supplementary Table 9:** Comparison of tau measurements in between subject groups

**Supplementary Table 10:** UFM1 and UFSP2 protein correlations with DDR gene expression in AD patient temporal cortex

**Supplementary Table 11:** UFM1 and UFSP2 protein correlations with seven UPR genes expression in AD patient temporal cortex

### SFig 1: UFMylation gene expression is dysregulated in AD excitatory neurons

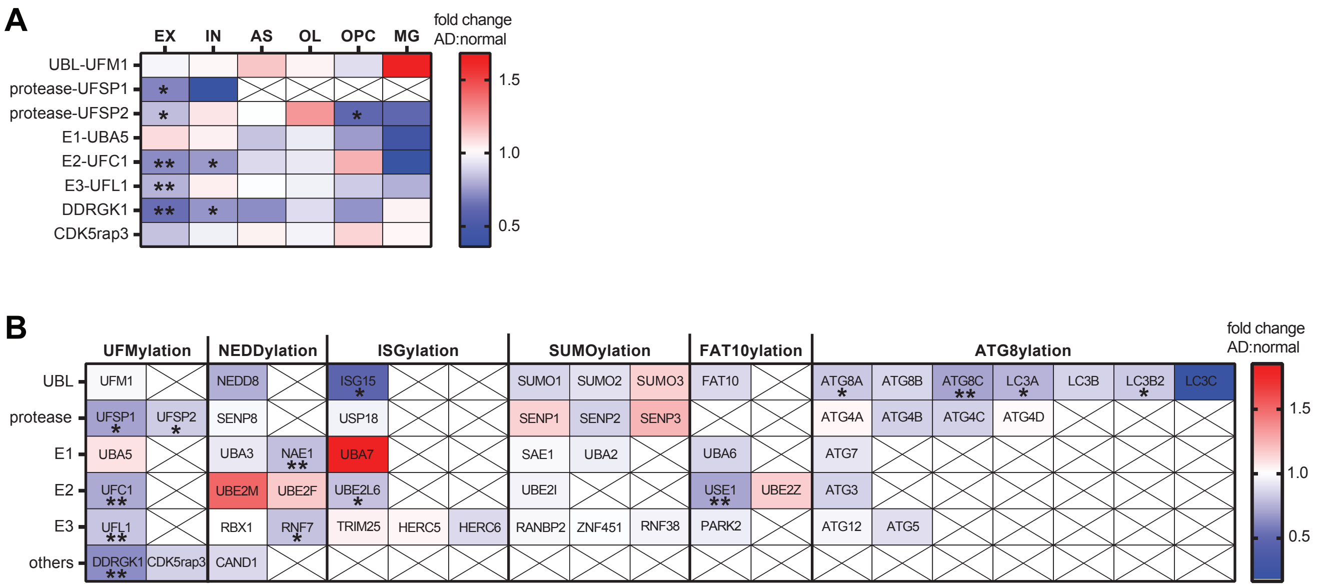

### SFig 2: Optimization and assessments of the MSD ELISA for UFM1

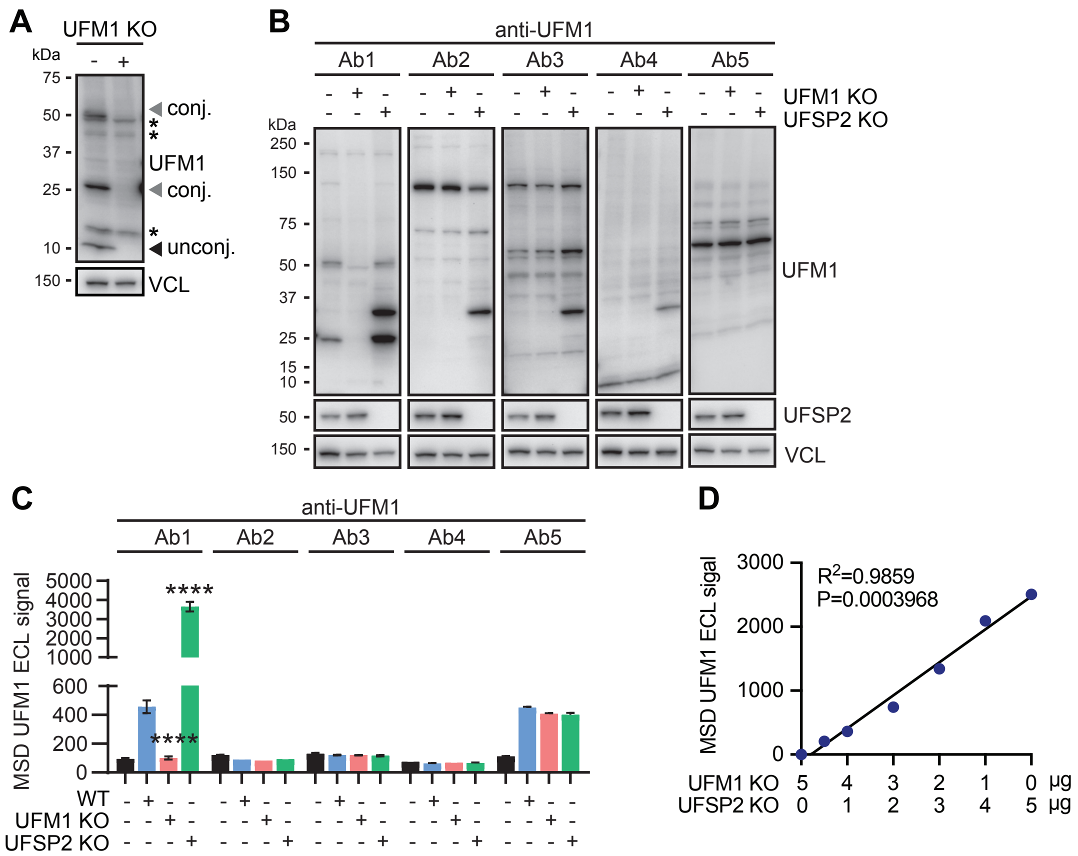

### SFig 3: Assessments of MSD ELISA of UFSP2

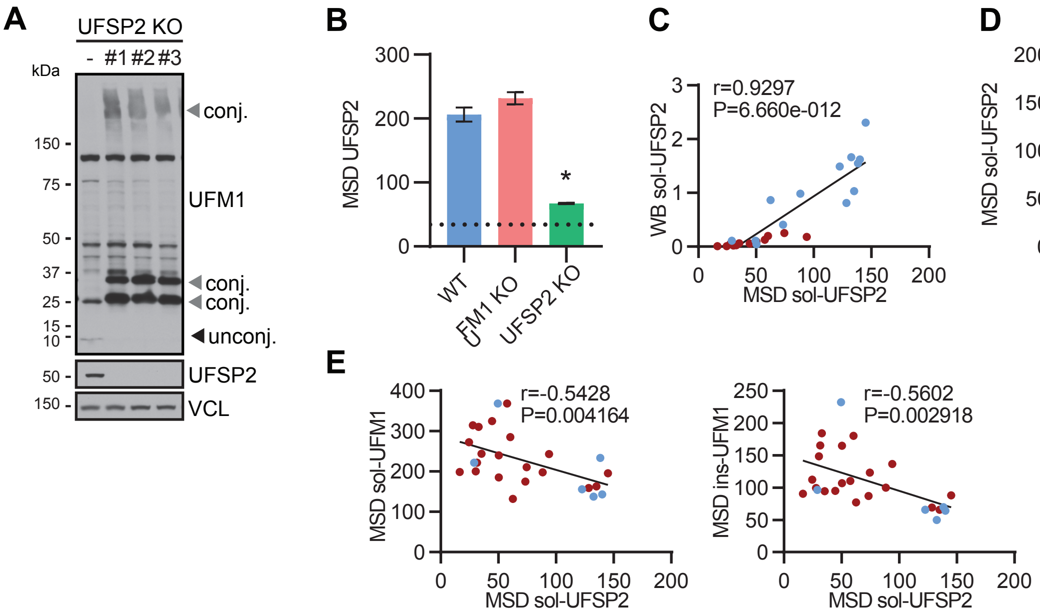

### SFig 4: Correlations of UFM1 and UFSP2 with biochemical AD measures from the temporal cortex

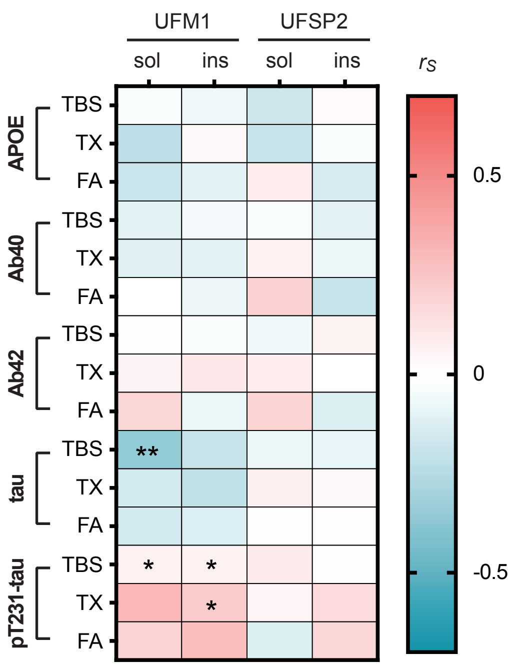

### SFig 5: UFSP2 KO does not affect susceptibility of neuroprogenitor cells towards DNA damage

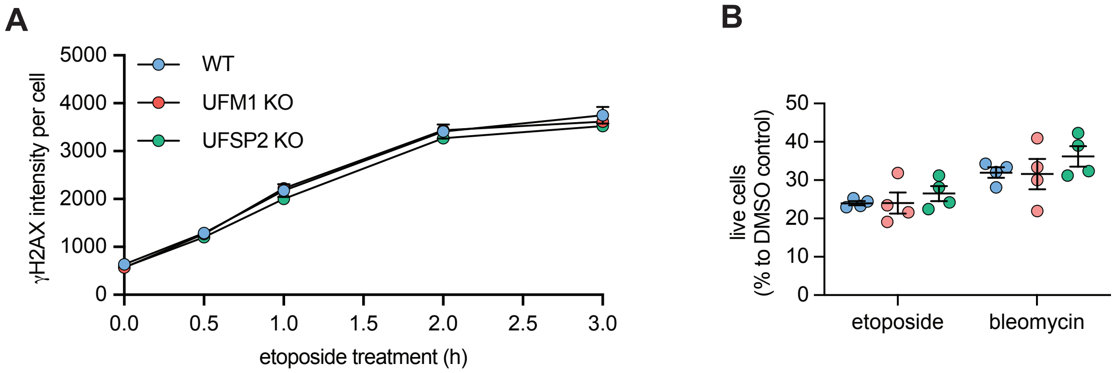

**Supplemental Table 1: Subject characteristics of exploratory cohort.**

| **Cohort** | **N** | **Male** (%) | **age** median (min, max) | **disease duration** median (min, max) | **Braak tangle stage** median (min, max) | **Thal amyloid phase** median (min, max) |
|---|---|---|---|---|---|---|
| control | 13 | 7 (53.8%) | 73 (63, 91) | NA | 0 (0, 3) | 1 (0, 1) |
| AD | 13 | 6 (46.2%) | 80 (64, 89) | 8 (7, 16) | 6 (5, 6) | 5 (5, 5) |

AD=Alzheimer’s disease. Age and neuropathological scores are presented as median (min, max).

**Supplemental Table 2: Subject characteristics**

| | AD patients (N=72) | | Controls (N=41) | | |
|---|---|---|---|---|---|
| Variable | N | Median (minimum, maximum) or No. (%) of patients | N | Median (minimum, maximum) or No. (%) of patients | P-value |
| Age (years) | 72 | 82.3 (57.8, 95.9) | 41 | 85.5 (70.5, 94.7) | 0.1220 |
| Sex (Male) | 72 | 37 (51.4%) | 41 | 22 (53.7%) | 0.8469 |
| *APOE* genotype | 72 | | 22 | | N/A |
| ε2/ε3 | | 3 (4.2%) | | 1 (4.5%) | |
| ε2/ε4 | | 2 (2.8%) | | 0 (0.0%) | |
| ε3/ε3 | | 24 (33.3%) | | 17 (77.3%) | |
| ε3/ε4 | | 32 (44.4%) | | 4 (18.2%) | |
| ε4/ε4 | | 11 (15.3%) | | 0 (0.0%) | |
| Disease duration (years) | 71 | 8.8 (2.4, 23.8) | 0 | N/A | N/A |
| MMSE score | 61 | 13 (0, 27) | 0 | N/A | N/A |
| Braak stage | 72 | | 41 | | N/A |
| 0 | | 0 (0.0%) | | 1 (2.4%) | |
| I | | 0 (0.0%) | | 5 (12.2%) | |
| II | | 0 (0.0%) | | 8 (19.5%) | |
| III | | 0 (0.0%) | | 27 (65.9%) | |
| IV | | 17 (23.6%) | | 0 (0.0%) | |
| V | | 33 (45.8%) | | 0 (0.0%) | |
| VI | | 22 (30.6%) | | 0 (0.0%) | |
| Thal phase | 72 | | 41 | | N/A |
| 0 | | 0 (0.0%) | | 19 (46.3%) | |
| 1 | | 0 (0.0%) | | 4 (9.8%) | |
| 2 | | 0 (0.0%) | | 8 (19.5%) | |
| 3 | | 6 (8.3%) | | 10 (24.4%) | |
| 4 | | 9 (12.5%) | | 0 (0.0%) | |
| 5 | | 57 (79.2%) | | 0 (0.0%) | |
| Biochemical data (temporal cortex) | 72 | | | | |
| apoE TBS | | 480.7 (7.8, 1244.4) | | N/A | N/A |
| apoE TX | | 243.4 (33.8, 524.3) | | N/A | N/A |
| apoE FA | | 330.8 (47.4, 1499.2) | | N/A | N/A |
| Aβ40 TBS | | 69.8 (0.0, 8212.2) | | N/A | N/A |
| Aβ40 TX | | 271.8 (8.2, 9482.8) | | N/A | N/A |
| Aβ40 FA | | 964.7 (68.1, 186998.4) | | N/A | N/A |
| Aβ42 TBS | | 631.2 (174.7, 2800.6) | | N/A | N/A |
| Aβ42 TX | | 1511.6 (363.8, 6234.5) | | N/A | N/A |
| Aβ42 FA | | 57.9 (12.8, 220.2) | | N/A | N/A |
| tau TBS | | 4.7 (0.1, 10.6) | | N/A | N/A |
| tau TX | | 1488.0 (63.1, 4134.1) | | N/A | N/A |
| tau FA | | 11.7 (1.5, 27.1) | | N/A | N/A |
| pT231-tau TBS | | 10.5 (1.9, 26.1) | | N/A | N/A |
| pT231-tau TX | | 5.4 (1.3, 16.9) | | N/A | N/A |
| pT231-tau FA | | 1404.9 (57.0, 11311.7) | | N/A | N/A |

AD=Alzheimer’s disease; MMSE=Mini mental state examination; TBS, TX, FA represent different sequential fractions. P-values result from a Wilcoxon rank sum test (continuous and ordinal variables) or Fisher’s exact test (categorical variables).

**Supplemental Table 3: Transformation of variables for linear regression analyses**

| Variable | Transformation |
|---|---|
| Age | None |
| Disease duration | None |
| MMSE score | None |
| Temporal cortex | |
| sol total tau | None |
| ins total tau | Square root |
| sol p396/404-tau | Square root |
| ins p396/404-tau | Cube root |
| sol UFSP2 | Square root |
| ins UFSP2 | None |
| sol UFM1 | None |
| ins UFM1 | None |
| Frontal cortex | |
| sol total tau | None |
| ins total tau | Square root |
| sol p396/404-tau | Cube root |
| ins p396/404-tau | Square root |
| sol UFSP2 | Square root |
| ins UFSP2 | None |
| sol UFM1 | None |
| ins UFM1 | None |

MMSE=mini mental state examination; sol=soluble; ins=insoluble.

### Supplemental Table 4: Differential single nuclei expression of UBL pathway components in samples with and without AD pathology

| | | | no-AD pathology vs AD pathology differential expression | | | | | | | |
|---|---|---|---|---|---|---|---|---|---|---|
| Cell type | Role of genes | Genes | IndModel .adj.pvals | no.pathology. mean | pathology. mean | FC | MixedModel. z | MixedModel. p | DEGs.In d.Model | DEGs.Ind.Mix. models |
| EX | UBL | UFM1 | 2.07E-26 | 0.1666 | 0.1620 | 0.9724 | -0.4130 | 0.6796 | FALSE | FALSE |
| IN | UBL | UFM1 | 1.26E-02 | 0.1057 | 0.1080 | 1.0220 | -0.1872 | 0.8515 | FALSE | FALSE |
| AS | UBL | UFM1 | 5.34E-01 | 0.0344 | 0.0394 | 1.1445 | -0.7531 | 0.4514 | FALSE | FALSE |
| OL | UBL | UFM1 | 6.46E-01 | 0.0409 | 0.0423 | 1.0347 | 0.3771 | 0.7061 | FALSE | FALSE |
| OPC | UBL | UFM1 | 8.57E-01 | 0.0550 | 0.0504 | 0.9159 | -0.3088 | 0.7574 | FALSE | FALSE |
| MG | UBL | UFM1 | 7.60E-01 | 0.0196 | 0.0330 | 1.6818 | 0.6719 | 0.5016 | FALSE | FALSE |
| EX | UFM1 Protease | UFSP1 | 7.18E-03 | 0.0031 | 0.0022 | 0.7053 | NA | NA | **TRUE** | FALSE |
| IN | UFM1 Protease | UFSP1 | 1.85E-01 | 0.0014 | 0.0005 | 0.3602 | NA | NA | FALSE | FALSE |
| AS | UFM1 Protease | UFSP1 | N/A | | | | | | | |
| OL | UFM1 Protease | UFSP1 | N/A | | | | | | | |
| OPC | UFM1 Protease | UFSP1 | N/A | | | | | | | |
| MG | UFM1 Protease | UFSP1 | N/A | | | | | | | |
| EX | UFM1 Protease | UFSP2 | 3.56E-16 | 0.0669 | 0.0559 | 0.8347 | -0.0223 | 0.9822 | **TRUE** | **FALSE** |
| IN | UFM1 Protease | UFSP2 | 2.24E-01 | 0.0615 | 0.0655 | 1.0654 | 0.0434 | 0.9654 | FALSE | FALSE |
| AS | UFM1 Protease | UFSP2 | 9.49E-01 | 0.0492 | 0.0489 | 0.9938 | 0.7409 | 0.4587 | FALSE | FALSE |
| OL | UFM1 Protease | UFSP2 | 7.69E-03 | 0.0354 | 0.0450 | 1.2706 | 2.7633 | 0.0057 | **TRUE** | FALSE |
| OPC | UFM1 Protease | UFSP2 | 6.15E-01 | 0.0186 | 0.0113 | 0.6112 | -1.6260 | 0.1039 | FALSE | FALSE |
| MG | UFM1 Protease | UFSP2 | 6.15E-01 | 0.0186 | 0.0113 | 0.6112 | -1.6260 | 0.1039 | FALSE | FALSE |
| EX | UFM1 E1 | UBA5 | 1.76E-20 | 0.1002 | 0.1095 | 1.0930 | 1.4667 | 0.1424 | FALSE | FALSE |
| IN | UFM1 E1 | UBA5 | 7.18E-02 | 0.0565 | 0.0588 | 1.0413 | -0.1776 | 0.8591 | FALSE | FALSE |
| AS | UFM1 E1 | UBA5 | 2.65E-01 | 0.0529 | 0.0452 | 0.8542 | -1.6192 | 0.1054 | FALSE | FALSE |
| OL | UFM1 E1 | UBA5 | 9.36E-01 | 0.0150 | 0.0142 | 0.9483 | -0.2372 | 0.8125 | FALSE | FALSE |
| OPC | UFM1 E1 | UBA5 | 5.74E-01 | 0.0566 | 0.0429 | 0.7584 | -1.3782 | 0.1681 | FALSE | FALSE |
| MG | UFM1 E1 | UBA5 | 9.39E-01 | 0.0121 | 0.0053 | 0.4386 | -2.2390 | 0.0252 | FALSE | FALSE |
| EX | UFM1 E2 | UFC1 | 1.63E-75 | 0.1422 | 0.1026 | 0.7220 | -5.5931 | 0.0000 | **TRUE** | **TRUE** |
| IN | UFM1 E2 | UFC1 | 3.06E-06 | 0.0618 | 0.0465 | 0.7525 | -2.4667 | 0.0136 | **TRUE** | FALSE |
| AS | UFM1 E2 | UFC1 | 6.49E-01 | 0.0521 | 0.0472 | 0.9057 | -0.1046 | 0.9167 | FALSE | FALSE |
| OL | UFM1 E2 | UFC1 | 8.94E-01 | 0.0314 | 0.0295 | 0.9413 | 0.4133 | 0.6794 | FALSE | FALSE |
| OPC | UFM1 E2 | UFC1 | 9.29E-01 | 0.0437 | 0.0526 | 1.2021 | 0.8315 | 0.4057 | FALSE | FALSE |
| MG | UFM1 E2 | UFC1 | 1.66E-01 | 0.0358 | 0.0131 | 0.3665 | -2.6684 | 0.0076 | FALSE | FALSE |
| EX | UFM1 E3 | UFL1 | 3.59E-57 | 0.1418 | 0.1156 | 0.8155 | -4.7652 | 0.0000 | **TRUE** | **TRUE** |
| IN | UFM1 E3 | UFL1 | 7.45E-03 | 0.1058 | 0.1107 | 1.0464 | -0.0663 | 0.9471 | FALSE | FALSE |

| | | | | | | | | | | |
|---|---|---|---|---|---|---|---|---|---|---|
| AS | UFM1 E3 | UFL1 | 5.50E-01 | 0.1507 | 0.1494 | 0.9914 | -0.1384 | 0.8899 | FALSE | FALSE |
| OL | UFM1 E3 | UFL1 | 4.22E-01 | 0.0957 | 0.0918 | 0.9590 | -1.1643 | 0.2443 | FALSE | FALSE |
| OPC | UFM1 E3 | UFL1 | 6.15E-01 | 0.1531 | 0.1331 | 0.8697 | -0.5380 | 0.5906 | FALSE | FALSE |
| MG | UFM1 E3 | UFL1 | 9.68E-01 | 0.0439 | 0.0355 | 0.8081 | -0.4989 | 0.6179 | FALSE | FALSE |
| EX | UFM1 E3 complex | DDRGK1 | 3.40E-140 | 0.2460 | 0.1580 | 0.6422 | -6.8151 | 0.0000 | **TRUE** | **TRUE** |
| IN | UFM1 E3 complex | DDRGK1 | 1.87E-07 | 0.0768 | 0.0571 | 0.7431 | -1.0690 | 0.2851 | **TRUE** | FALSE |
| AS | UFM1 E3 complex | DDRGK1 | 6.18E-02 | 0.0930 | 0.0675 | 0.7256 | -0.8582 | 0.3908 | FALSE | FALSE |
| OL | UFM1 E3 complex | DDRGK1 | 6.66E-01 | 0.0619 | 0.0566 | 0.9146 | -1.5869 | 0.1125 | FALSE | FALSE |
| OPC | UFM1 E3 complex | DDRGK1 | 7.25E-01 | 0.0707 | 0.0522 | 0.7385 | -0.4086 | 0.6829 | FALSE | FALSE |
| MG | UFM1 E3 complex | DDRGK1 | 9.11E-01 | 0.0125 | 0.0129 | 1.0332 | 0.2798 | 0.7797 | FALSE | FALSE |
| EX | UFM1 E3 complex | CDK5RAP3 | 2.87E-57 | 0.1903 | 0.1619 | 0.8508 | -1.1861 | 0.2356 | FALSE | FALSE |
| IN | UFM1 E3 complex | CDK5RAP3 | 4.18E-03 | 0.1078 | 0.1031 | 0.9566 | 0.5762 | 0.5645 | FALSE | FALSE |
| AS | UFM1 E3 complex | CDK5RAP3 | 7.78E-01 | 0.1971 | 0.2048 | 1.0393 | 0.2635 | 0.7922 | FALSE | FALSE |
| OL | UFM1 E3 complex | CDK5RAP3 | 8.25E-01 | 0.1124 | 0.1085 | 0.9649 | -0.6207 | 0.5348 | FALSE | FALSE |
| OPC | UFM1 E3 complex | CDK5RAP3 | 9.54E-01 | 0.1393 | 0.1545 | 1.1096 | 1.5649 | 0.1176 | FALSE | FALSE |
| MG | UFM1 E3 complex | CDK5RAP3 | 8.81E-01 | 0.0812 | 0.0833 | 1.0251 | 0.3773 | 0.7060 | FALSE | FALSE |
| EX | UBL | ISG15 | 6.95E-50 | 0.0531 | 0.0253 | 0.4759 | -3.3039 | 0.0010 | **TRUE** | FALSE |
| EX | UBL | NEDD8 | 1.71E-01 | 0.0007 | 0.0005 | 0.7471 | NA | NA | FALSE | FALSE |
| EX | UBL | SUMO1 | 6.66E-71 | 0.3539 | 0.3060 | 0.8647 | -2.4086 | 0.0160 | FALSE | FALSE |
| EX | UBL | SUMO2 | 2.11E-22 | 0.0626 | 0.0566 | 0.9033 | -1.8748 | 0.0608 | FALSE | FALSE |
| EX | UBL | SUMO3 | 3.66E-15 | 0.2843 | 0.3279 | 1.1534 | 3.3783 | 0.0007 | FALSE | FALSE |
| EX | UBL | FAT10 | 2.06E-91 | 0.5297 | 0.4623 | 0.8728 | -3.0750 | 0.0021 | FALSE | FALSE |
| EX | UBL (ATG8) | GABARAP | 1.91E-06 | 0.0287 | 0.0235 | 0.8218 | 0.2695 | 0.7876 | **TRUE** | FALSE |
| EX | UBL (ATG8) | GABARAPL1 | 3.53E-90 | 0.5040 | 0.4369 | 0.8670 | -3.5538 | 0.0004 | FALSE | FALSE |
| EX | UBL (ATG8) | GABARAPL2 | 1.79E-145 | 0.5547 | 0.3918 | 0.7063 | -4.9819 | 0.0000 | **TRUE** | **TRUE** |
| EX | UBL (ATG8) | MAP1LC3A | 1.48E-100 | 0.3627 | 0.2764 | 0.7622 | -3.2186 | 0.0013 | **TRUE** | FALSE |
| EX | UBL (ATG8) | MAP1LC3B | 5.98E-68 | 0.3353 | 0.2876 | 0.8577 | -4.7490 | 0.0000 | FALSE | FALSE |
| EX | UBL (ATG8) | MAP1LC3B2 | 6.48E-23 | 0.0440 | 0.0362 | 0.8234 | -3.9275 | 0.0001 | **TRUE** | FALSE |
| EX | NEDD8 Protease | SENP8 | 3.63E-08 | 0.0314 | 0.0304 | 0.9679 | 0.8687 | 0.3850 | FALSE | FALSE |
| EX | ISG15 Protease | USP18 | 4.44E-07 | 0.0137 | 0.0131 | 0.9536 | -0.8783 | 0.3798 | FALSE | FALSE |
| EX | SUMO Protease | SENP1 | 1.11E-08 | 0.0632 | 0.0723 | 1.1444 | 2.9842 | 0.0028 | FALSE | FALSE |
| EX | SUMO Protease | SENP2 | 5.56E-69 | 0.2796 | 0.2380 | 0.8514 | -1.7946 | 0.0727 | FALSE | FALSE |
| EX | SUMO Protease | SENP3 | 8.36E-01 | 0.0002 | 0.0002 | 1.2371 | NA | NA | FALSE | FALSE |
| EX | SUMO Protease | SENP5 | 3.59E-66 | 0.3714 | 0.3432 | 0.9240 | -1.6980 | 0.0895 | FALSE | FALSE |
| EX | SUMO Protease | SENP6 | 1.30E-49 | 0.5918 | 0.5936 | 1.0031 | 1.9740 | 0.0484 | FALSE | FALSE |
| EX | SUMO Protease | SENP7 | 3.60E-38 | 0.2050 | 0.1996 | 0.9734 | -0.2334 | 0.8155 | FALSE | FALSE |
| EX | ATG8 Protease | ATG4A | 8.58E-06 | 0.0256 | 0.0265 | 1.0318 | 0.4072 | 0.6839 | FALSE | FALSE |
| EX | ATG8 Protease | ATG4B | 1.62E-105 | 0.3164 | 0.2696 | 0.8521 | -2.2549 | 0.0241 | FALSE | FALSE |
| EX | ATG8 Protease | ATG4C | 5.83E-64 | 0.1667 | 0.1420 | 0.8520 | -4.4933 | 0.0000 | FALSE | FALSE |

| | | | | | | | | | | |
|---|---|---|---|---|---|---|---|---|---|---|
| EX | ATG8 Protease | ATG4D | 4.51E-22 | 0.0818 | 0.0833 | 1.0189 | -0.5735 | 0.5663 | FALSE | FALSE |
| EX | ISG15 E1 | UBA7 | 3.69E-01 | 0.0009 | 0.0017 | 1.8508 | NA | NA | FALSE | FALSE |
| EX | NEDD8 E1 | UBA3 | 9.15E-56 | 0.2511 | 0.2318 | 0.9230 | -0.6925 | 0.4886 | FALSE | FALSE |
| EX | NEDD8 E1 | NAE1 | 7.10E-90 | 0.2427 | 0.1933 | 0.7967 | -4.7057 | 0.0000 | **TRUE** | **TRUE** |
| EX | SUMO E1 | SAE1 | 3.52E-43 | 0.2413 | 0.2350 | 0.9741 | -0.0079 | 0.9937 | FALSE | FALSE |
| EX | SUMO E1 | UBA2 | 8.91E-62 | 0.3508 | 0.3298 | 0.9400 | -0.3340 | 0.7383 | FALSE | FALSE |
| EX | ATG8 E1 | ATG7 | 5.75E-102 | 0.8162 | 0.7454 | 0.9132 | -4.6332 | 0.0000 | FALSE | FALSE |
| EX | ISG15 E2 | UBE2L6 | 1.88E-21 | 0.0510 | 0.0409 | 0.8018 | -2.5585 | 0.0105 | **TRUE** | FALSE |
| EX | NEDD8 E2 | UBE2M | 7.81E-02 | 0.2491 | 0.3755 | 1.5076 | 3.3878 | 0.0007 | FALSE | FALSE |
| EX | NEDD8 E2 | UBE2F-SCLY | 1.38E-06 | 0.0245 | 0.0287 | 1.1719 | 0.3064 | 0.7593 | FALSE | FALSE |
| EX | FAT10 E2 | USE1 | 3.30E-44 | 0.0841 | 0.0600 | 0.7141 | -4.9904 | 0.0000 | **TRUE** | **TRUE** |
| EX | FAT10 E2 | UBE2Z | 1.64E-21 | 0.2347 | 0.2725 | 1.1610 | 2.5947 | 0.0095 | FALSE | FALSE |
| EX | ATG12 E2 | ATG10 | 5.47E-104 | 0.2630 | 0.2049 | 0.7790 | -2.5290 | 0.0114 | **TRUE** | FALSE |
| EX | SUMO E2 | UBE2I | 2.85E-44 | 0.2413 | 0.2305 | 0.9552 | 0.1179 | 0.9061 | FALSE | FALSE |
| EX | ATG8 E2 | ATG3 | 2.30E-17 | 0.0641 | 0.0548 | 0.8547 | -0.8928 | 0.3719 | FALSE | FALSE |
| EX | SUMO E3 | RNF38 | 1.78E-47 | 0.3094 | 0.2962 | 0.9572 | 0.0675 | 0.9461 | FALSE | FALSE |
| EX | ISG15 E3 | TRIM25 | 8.11E-04 | 0.0328 | 0.0345 | 1.0510 | 2.0130 | 0.0441 | FALSE | FALSE |
| EX | ISG15 E3 | HERC5 | 4.75E-01 | 0.0025 | 0.0026 | 1.0354 | NA | NA | FALSE | FALSE |
| EX | NEDD8 E3 | RBX1 | 4.81E-50 | 0.4145 | 0.4160 | 1.0036 | 0.5073 | 0.6119 | FALSE | FALSE |
| EX | NEDD8 E3 | RNF7 | 2.29E-46 | 0.1320 | 0.1065 | 0.8068 | -3.2946 | 0.0010 | **TRUE** | FALSE |
| EX | SUMO E3 | RANBP2 | 4.20E-65 | 0.4989 | 0.4845 | 0.9711 | -1.4443 | 0.1487 | FALSE | FALSE |
| EX | SUMO E3 | ZNF451 | 1.37E-52 | 0.4200 | 0.4146 | 0.9872 | 0.8150 | 0.4151 | FALSE | FALSE |
| EX | SUMO E3 | RNF38 | 1.78E-47 | 0.3094 | 0.2962 | 0.9572 | 0.0675 | 0.9461 | FALSE | FALSE |
| EX | FAT10 E3 | PARK2 | 2.47E-75 | 1.1431 | 1.0713 | 0.9372 | -3.0734 | 0.0021 | FALSE | FALSE |
| EX | ATG8 E3 | ATG12 | 4.48E-37 | 0.2013 | 0.1935 | 0.9609 | -0.1696 | 0.8653 | FALSE | FALSE |
| EX | ATG8 E3 | ATG5 | 2.34E-62 | 0.2100 | 0.1885 | 0.8976 | -4.9821 | 0.0000 | FALSE | FALSE |

Meta analysis of published single cell transcriptome data [35]. EX=excitatory neurons, IN=inhibitory neurons, AS=astrocytes, OL=oligodendrocytes, OPC=oligodendrocyte precursor cells, MG=microglia, UBL=ubiquitin-like protein. IndModel.adj.pvals= fdr-adjusted p-values, two-sided 2-sided Wilcoxon-rank-sum test, FC=fold change of AD pathology mean value relative to no-AD pathology mean value, MixedModel.z=z-score of Poisson mixed model, MixedModel.p=P-value of Poisson mixed model, DEGs.Ind.Model=logical indication of whether a gene meets the criteria fdr-adjusted P-value<0.01 and absolute log2 fold change > 0.25, DEGs.Ind.Mix.models=logical indication of whether a gene meets the criteria fdr-adjusted MixedModel.P<0.05 (frd-adjustment over genes meeting criteria in DEGs.Ind.Model)

**Supplemental Table 5: Correlation of UFM1 values with other UFMylation proteins in exploratory cohort**

| | $r_s$ (P-value) | | | | | | | |
|---|---|---|---|---|---|---|---|---|
| Variable | free UFM1 | UFSP1 | UFSP2 | UBA5 | UFC1 | UFL1 | DDRGK1 | CDK5RAP3 |
| total sol UFM1 | 0.14<br>(0.4901) | 0.01<br>(0.9709) | **-0.56**<br>**(0.0028)** | -0.33<br>(0.0956) | 0.14<br>(0.485) | -0.43<br>(0.028) | -0.12<br>(0.5435) | -0.36<br>(0.0668) |
| total ins UFM1 | -0.31<br>(0.121) | -0.24<br>(0.228) | **-0.66**<br>**(0.0002)** | -0.48<br>(0.0142) | 0.22<br>(0.2699) | -0.27<br>(0.1808) | 0.03<br>(0.8827) | -0.41<br>(0.0395) |

rs=Spearman correlation coefficient; sol=soluble, ins=insoluble. Total UFM1 was determined by MSD ELISA in the RIPA-soluble and -insoluble fraction in frontal cortex tissue of the exploratory cohort (all subjects combined). Other variables were determined by western blot of the soluble fraction. P-values < 0.0055 were considered statistically significant after applying a Bonferroni correction for multiple testing. Significant associations are shown in bold.

### Supplemental Table 6: Correlation of UFSP2 with UFM1 in human brain

| | **Association with sol UFSP2** | | | | **Association with ins UFSP2** | | | |
|---|---|---|---|---|---|---|---|---|
| | unadjusted analysis | | multivariable analysis | | unadjusted analysis | | multivariable analysis | |
| Cortex/Group/Variable | β (95% CI) | P-value | β (95% CI) | P-value | β (95% CI) | P-value | β (95% CI) | P-value |
| **Temporal Cortex** | | | | | | | | |
| Controls (N=41) | | | | | | | | |
| sol UFSP2 | N/A | N/A | N/A | N/A | -0.71 (-1.60, 0.18) | 0.1128 | -0.76 (-1.71, 0.18) | 0.1108 |
| ins UFSP2 | -0.71 (-1.60, 0.18) | 0.1128 | -0.76 (-1.71, 0.18) | 0.1108 | N/A | N/A | N/A | N/A |
| sol UFM1 | -0.01 (-1.03, 1.01) | 0.9808 | -0.03 (-1.10, 1.05) | 0.9603 | 9.79 (-2.93, 22.50) | 0.1276 | 8.60 (-4.74, 21.93) | 0.1993 |
| ins UFM1 | -0.93 (-1.49, -0.36) | 0.0019 | **-0.94 (-1.59, -0.29)** | **0.0058** | 2.27 (-5.89, 10.42) | 0.5776 | 4.47 (-4.59, 13.52) | 0.3238 |
| AD patients (N=72) | | | | | | | | |
| sol UFSP2 | N/A | N/A | N/A | N/A | -1.03 (-1.46, -0.59) | <0.0001 | **-1.01 (-1.46, -0.57)** | **<0.0001** |
| ins UFSP2 | -1.03 (-1.46, -0.59) | <0.0001 | **-1.01 (-1.46, -0.57)** | **<0.0001** | N/A | N/A | N/A | N/A |
| sol UFM1 | 0.24 (-0.24, 0.72) | 0.3304 | 0.19 (-0.31, 0.69) | 0.4435 | 11.73 (3.78, 19.69) | 0.0044 | **12.96 (4.69, 21.23)** | **0.0026** |
| ins UFM1 | -1.25 (-1.81, -0.69) | <0.0001 | **-1.24 (-1.84, -0.64)** | **0.0001** | 16.56 (6.28, 26.83) | 0.0020 | **18.45 (7.52, 29.37)** | **0.0013** |
| All subjects (N=113) | | | | | | | | |
| sol UFSP2 | N/A | N/A | N/A | N/A | -0.96 (-1.35, -0.58) | <0.0001 | **-0.96 (-1.36, -0.57)** | **<0.0001** |
| ins UFSP2 | -0.96 (-1.35, -0.58) | <0.0001 | **-0.96 (-1.36, -0.57)** | **<0.0001** | N/A | N/A | N/A | N/A |
| sol UFM1 | 0.19 (-0.24, 0.62) | 0.3822 | 0.15 (-0.29, 0.60) | 0.4995 | 11.17 (4.57, 17.77) | 0.0009 | **11.72 (4.86, 18.59)** | **0.0008** |
| ins UFM1 | -1.09 (-1.47, -0.70) | <0.0001 | **-1.10 (-1.53, -0.67)** | **<0.0001** | 9.05 (-4.94, 23.04) | 0.2047 | 11.10 (-2.59, 24.78) | 0.1119 |
| **Frontal Cortex** | | | | | | | | |
| Controls (N=41) | | | | | | | | |
| sol UFSP2 | N/A | N/A | N/A | N/A | -1.37 (-2.23, -0.52) | 0.0024 | **-1.85 (-2.83, -0.86)** | **0.0005** |
| ins UFSP2 | -1.37 (-2.23, -0.52) | 0.0024 | **-1.85 (-2.83, -0.86)** | **0.0005** | N/A | N/A | N/A | N/A |
| sol UFM1 | -1.03 (-2.36, 0.30) | 0.1266 | -1.13 (-2.58, 0.32) | 0.1213 | 26.93 (8.04, 45.83) | 0.0064 | 19.89 (1.23, 38.54) | 0.0373 |
| ins UFM1 | -1.73 (-2.54, -0.93) | <0.0001 | **-1.95 (-2.83, -1.06)** | **<0.0001** | 9.38 (-5.20, 23.96) | 0.2007 | 12.02 (-2.11, 26.14) | 0.0929 |
| AD patients (N=72) | | | | | | | | |
| sol UFSP2 | N/A | N/A | N/A | N/A | -1.46 (-2.18, -0.74) | 0.0001 | **-1.46 (-2.24, -0.69)** | **0.0003** |
| ins UFSP2 | -1.46 (-2.18, -0.74) | 0.0001 | **-1.46 (-2.24, -0.69)** | **0.0003** | N/A | N/A | N/A | N/A |
| sol UFM1 | -0.61 (-1.28, 0.06) | 0.0734 | -0.62 (-1.34, 0.10) | 0.0892 | 12.01 (3.24, 20.79) | 0.0080 | **11.45 (2.24, 20.67)** | **0.0156** |
| ins UFM1 | -1.47 (-2.25, -0.69) | 0.0003 | **-1.51 (-2.37, -0.65)** | **0.0008** | 18.71 (8.17, 29.26) | 0.0007 | **17.96 (6.55, 29.36)** | **0.0025** |
| All subjects (N=113) | | | | | | | | |
| sol UFSP2 | N/A | N/A | N/A | N/A | -1.42 (-1.96, -0.89) | <0.0001 | **-1.61 (-2.21, -1.02)** | **<0.0001** |
| ins UFSP2 | -1.42 (-1.96, -0.89) | <0.0001 | **-1.61 (-2.21, -1.02)** | **<0.0001** | N/A | N/A | N/A | N/A |
| sol UFM1 | -0.70 (-1.28, -0.11) | 0.0199 | -0.73 (-1.36, -0.10) | 0.0240 | 17.20 (3.27, 31.12) | 0.0155 | **13.15 (5.07, 21.23)** | **0.0014** |
| ins UFM1 | -1.60 (-2.14, -1.05) | <0.0001 | **-1.73 (-2.33, -1.13)** | **<0.0001** | 15.33 (6.55, 24.12) | 0.0006 | **15.57 (6.92, 24.22)** | **0.0004** |

sol=soluble; ins=insoluble; β=regression coefficient; CI=confidence intervals; AD=Alzheimer's disease. β values, 95% CIs, and P-values for the separate control and AD groups result from linear regression models. β values are interpreted as the increase in mean UFSP2 (on the square root scale when examining sol UFSP2) corresponding to 1 SD increase for the shown continuous variables, which were examined untransformed, or square root transformed (sol UFSP2). Multivariable models for controls were adjusted for age, sex, Braak stage, and Thal phase, and multivariable models for AD patients were adjusted for age, sex, presence of *APOE* ε4, Braak stage, and Thal phase. β values, 95% CIs, and P-values for the analysis of all subjects results from a random effects meta-analysis combining the separate results from the control and AD groups. After applying a Bonferroni correction for multiple testing P-values < 0.0167 are considered as statistically significant. Significant findings from the multivariate analysis are shown in bold. Note: there was no significant association of soluble UFM1 with insoluble UFM1 in any of the models.

### Supplemental Table 7: Associations of primary clinical and disease parameters with UFSP2 and UFM1

| | Association with sol UFSP2 | | Association with ins UFSP2 | | Association with sol UFM1 | | Association with ins UFM1 | |
|---|---|---|---|---|---|---|---|---|
| Cortex/Group/Variable | β (95% CI) | P-value | β (95% CI) | P-value | β (95% CI) | P-value | β (95% CI) | P-value |
| **Temporal Cortex** | | | | | | | | |
| Controls (N=41) | | | | | | | | |
| Age (10 year increase) | 0.66 (-0.57, 1.88) | 0.2841 | 5.34 (-10.23, 20.90) | 0.4913 | 4.17 (-26.98, 35.32) | 0.7875 | -30.26 (-55.19, -5.32) | 0.0188 |
| Sex (Male) | -0.20 (-1.91, 1.52) | 0.8186 | 12.72 (-9.06, 34.50) | 0.2439 | 11.10 (-32.49, 54.69) | 0.6088 | -11.15 (-46.05, 23.74) | 0.5209 |
| Presence of *APOE* ε4 | -0.05 (-3.49, 3.38) | 0.9742 | -15.26 (-55.46, 24.93) | 0.4353 | -18.92 (-107.04, 69.20) | 0.6573 | 9.35 (-17.80, 36.50) | 0.4787 |
| Braak stage (1 unit increase) | 0.19 (-0.80, 1.17) | 0.7014 | -0.36 (-12.89, 12.16) | 0.9535 | 8.66 (-16.42, 33.73) | 0.4883 | -6.68 (-26.75, 13.39) | 0.5041 |
| Thal phase (1 unit increase) | -0.07 (-0.69, 0.56) | 0.8328 | 4.82 (-3.14, 12.77) | 0.2274 | 5.04 (-10.89, 20.96) | 0.5252 | -5.04 (-17.78, 7.71) | 0.4282 |
| AD patients (N=72) | | | | | | | | |
| Age (10 year increase) | 0.50 (-0.25, 1.25) | 0.1892 | -4.82 (-18.03, 8.39) | 0.4690 | 21.35 (-8.03, 50.74) | 0.1515 | -6.10 (-17.95, 5.75) | 0.3079 |
| Sex (Male) | -0.31 (-1.39, 0.76) | 0.5619 | 6.83 (-12.14, 25.80) | 0.4748 | -15.74 (-57.94, 26.47) | 0.4592 | -7.04 (-24.06, 9.98) | 0.4121 |
| Presence of *APOE* ε4 | -0.22 (-1.31, 0.86) | 0.6834 | 3.77 (-15.40, 22.94) | 0.6961 | 7.63 (-35.25, 50.51) | 0.7236 | 4.60 (-12.90, 22.09) | 0.6016 |
| Disease duration (5 year increase) | 0.58 (-0.15, 1.31) | 0.1164 | -2.26 (-15.42, 10.91) | 0.7332 | -7.03 (-36.25, 22.18) | 0.6322 | -9.38 (-20.88, 2.11) | 0.1077 |
| MMSE score (5 unit increase) | 0.08 (-0.44, 0.60) | 0.7553 | 1.89 (-7.28, 11.06) | 0.6810 | 3.22 (-16.50, 22.94) | 0.7446 | 4.19 (-3.61, 11.99) | 0.2864 |
| Braak stage (1 unit increase) | -0.22 (-1.02, 0.59) | 0.5899 | -4.09 (-18.26, 10.07) | 0.5659 | 5.13 (-26.37, 36.62) | 0.7464 | 9.54 (-3.18, 22.25) | 0.1392 |
| Thal phase (1 unit increase) | -0.06 (-0.98, 0.87) | 0.9059 | 3.21 (-13.15, 19.56) | 0.6966 | 15.45 (-20.92, 51.82) | 0.3996 | 1.22 (-13.46, 15.91) | 0.8684 |
| All subjects (N=113) | | | | | | | | |
| Age (10 year increase) | 0.54 (-0.08, 1.17) | 0.0890 | -0.49 (-10.33, 9.36) | 0.9229 | 13.13 (-7.70, 33.96) | 0.2166 | -15.78 (-38.98, 7.42) | 0.1826 |
| Sex (Male) | -0.28 (-1.17, 0.61) | 0.5380 | 9.42 (-4.53, 23.36) | 0.1857 | -2.54 (-32.08, 27.00) | 0.8660 | -7.85 (-22.82, 7.12) | 0.3042 |
| Presence of *APOE* ε4 | -0.21 (-1.22, 0.81) | 0.6900 | -0.06 (-16.89, 16.76) | 0.9940 | 2.11 (-35.37, 39.60) | 0.9121 | 6.10 (-8.12, 20.32) | 0.4008 |
| Braak stage (1 unit increase) | -0.05 (-0.66, 0.56) | 0.8646 | -1.97 (-11.10, 7.16) | 0.6723 | 7.31 (-11.76, 26.39) | 0.4524 | 3.20 (-12.31, 18.70) | 0.6861 |
| Thal phase (1 unit increase) | -0.06 (-0.57, 0.44) | 0.8082 | 4.52 (-2.42, 11.45) | 0.2017 | 6.67 (-7.46, 20.80) | 0.3551 | -2.39 (-11.76, 6.97) | 0.6162 |
| **Frontal Cortex** | | | | | | | | |
| Controls (N=41) | | | | | | | | |
| Age (10 year increase) | 0.69 (-1.06, 2.43) | 0.4321 | 12.72 (-10.47, 35.92) | 0.2732 | 8.29 (-15.09, 31.67) | 0.4768 | -9.93 (-37.86, 18.00) | 0.4754 |
| Sex (Male) | 0.02 (-2.43, 2.46) | 0.9889 | 14.29 (-18.16, 46.74) | 0.3777 | 12.63 (-20.08, 45.34) | 0.4387 | 0.76 (-38.32, 39.83) | 0.9688 |
| Presence of *APOE* ε4 | 0.13 (-3.96, 4.23) | 0.9472 | 19.31 (-33.42, 72.03) | 0.4518 | 6.69 (-48.69, 62.07) | 0.8025 | 40.17 (-16.96, 97.29) | 0.1569 |
| Braak stage (1 unit increase) | -0.09 (-1.50, 1.32) | 0.8970 | **25.32 (6.66, 43.99)** | **0.0092** | 11.71 (-7.10, 30.53) | 0.2149 | -11.07 (-33.54, 11.41) | 0.3246 |
| Thal phase (1 unit increase) | -0.04 (-0.93, 0.86) | 0.9358 | 7.37 (-4.48, 19.23) | 0.2152 | 2.05 (-9.90, 14.00) | 0.7300 | 14.58 (0.31, 28.86) | 0.0455 |
| AD patients (N=72) | | | | | | | | |
| Age (10 year increase) | 21.35 (-8.03, 50.74) | 0.1515 | -6.10 (-17.95, 5.75) | 0.3079 | -17.09 (-37.06, 2.87) | 0.0920 | -12.51 (-26.42, 1.40) | 0.0772 |
| Sex (Male) | -15.74 (-57.94, 26.47) | 0.4592 | -7.04 (-24.06, 9.98) | 0.4121 | -19.04 (-47.72, 9.63) | 0.1894 | -5.46 (-25.44, 14.52) | 0.5870 |
| Presence of *APOE* ε4 | 7.63 (-35.25, 50.51) | 0.7236 | 4.60 (-12.90, 22.09) | 0.6016 | 22.43 (-6.72, 51.57) | 0.1293 | 8.99 (-11.76, 29.74) | 0.3905 |
| Disease duration (5 year increase) | -7.03 (-36.25, 22.18) | 0.6322 | -9.38 (-20.88, 2.11) | 0.1077 | -3.89 (-23.74, 15.95) | 0.6964 | 3.62 (-10.21, 17.44) | 0.6029 |
| MMSE score (5 unit increase) | 3.22 (-16.50, 22.94) | 0.7446 | 4.19 (-3.61, 11.99) | 0.2864 | -3.06 (-14.44, 8.32) | 0.5923 | 7.60 (-1.61, 16.82) | 0.1039 |
| Braak stage (1 unit increase) | 5.13 (-26.37, 36.62) | 0.7464 | 9.54 (-3.18, 22.25) | 0.1392 | -10.63 (-32.39, 11.12) | 0.3329 | 3.07 (-11.90, 18.04) | 0.6833 |
| Thal phase (1 unit increase) | 15.45 (-20.92, 51.82) | 0.3996 | 1.22 (-13.46, 15.91) | 0.8684 | 9.55 (-15.57, 34.68) | 0.4504 | 15.45 (-1.83, 32.74) | 0.0788 |
| All subjects (N=113) | | | | | | | | |
| Age (10 year increase) | 13.13 (-7.70, 33.96) | 0.2166 | -15.78 (-38.98, 7.42) | 0.1826 | -5.05 (-29.89, 19.79) | 0.6902 | -11.98 (-24.17, 0.20) | 0.0539 |
| Sex (Male) | -2.54 (-32.08, 27.00) | 0.8660 | -7.85 (-22.82, 7.12) | 0.3042 | -4.06 (-35.05, 26.94) | 0.7975 | -4.14 (-21.55, 13.27) | 0.6410 |
| Presence of *APOE* ε4 | 2.11 (-35.37, 39.60) | 0.9121 | 6.10 (-8.12, 20.32) | 0.4008 | 18.73 (-6.31, 43.77) | 0.1426 | 14.46 (-8.78, 37.69) | 0.2227 |
| Braak stage (1 unit increase) | 7.31 (-11.76, 26.39) | 0.4524 | 3.20 (-12.31, 18.70) | 0.6861 | 1.27 (-20.58, 23.12) | 0.9091 | -1.64 (-14.71, 11.42) | 0.8052 |
| Thal phase (1 unit increase) | 6.67 (-7.46, 20.80) | 0.3551 | -2.39 (-11.76, 6.97) | 0.6162 | 3.40 (-7.06, 13.86) | 0.5242 | **14.93 (4.22, 25.63)** | **0.0063** |

sol=soluble; ins=insoluble; β=regression coefficient; CI=confidence intervals; AD=Alzheimer's disease. β values, 95% CIs, and P-values for the separate control and AD groups result from multivariate linear regression models. β values are interpreted as the increase in mean UFSP2 (on the square root scale when examining sol UFSP2) corresponding to presence of the given characteristic (categorical variables) or the increase given in parenthesis (continuous variables, which were examined on the untransformed, square root, cube root, or natural logarithm scale). Models for controls were adjusted for age, sex, Braak stage, and Thal phase, and models for AD patients were adjusted for age, sex, presence of *APOE* ε4, Braak stage, and Thal phase. β values, 95% CIs, and P-values for the analysis of all subjects results from a random effects meta-analysis combining the separate results from the control and AD groups. P-value < 0.01 (controls and all subjects) and < 0.0071 (AD patients) is considered as significant after applying a Bonferroni correction for multiple testing separately for each disease group and each outcome. Significant findings are shown in bold.

**Supplemental Table 8: Correlations of UFSP2 and UFM1 with common AD-related parameters in temporal cortex of AD patients**

| | $r_s$ (P-value) | | | |
|---|---|---|---|---|
| Variable Fraction | sol UFSP2 | ins UFSP2 | sol UFM1 | ins UFM1 |
| apoE TBS | -0.18 (0.1304) | 0.028 (0.8174) | -0.023 (0.8486) | -0.061 (0.6115) |
| apoE TX | -0.194 (0.1024) | -0.036 (0.7655) | -0.223 (0.0601) | 0.034 (0.7759) |
| apoE FA | 0.085 (0.4771) | -0.14 (0.2419) | -0.186 (0.1178) | -0.102 (0.3936) |
| Ab40 TBS | -0.029 (0.8108) | -0.099 (0.4081) | -0.102 (0.3929) | -0.049 (0.6799) |
| Ab40 TX | 0.07 (0.5573) | -0.074 (0.534) | -0.109 (0.3629) | -0.107 (0.372) |
| Ab40 FA | 0.202 (0.0889) | -0.193 (0.1042) | -0.003 (0.9822) | -0.073 (0.5402) |
| Ab42 TBS | -0.059 (0.6245) | 0.069 (0.5642) | -0.01 (0.9355) | -0.031 (0.7946) |
| Ab42 TX | 0.085 (0.4753) | -0.007 (0.9559) | 0.062 (0.6036) | 0.109 (0.3631) |
| Ab42 FA | 0.193 (0.1038) | -0.117 (0.3274) | 0.18 (0.1297) | -0.074 (0.535) |
| tau TBS | -0.078 (0.5166) | -0.087 (0.4658) | **-0.345 (0.003)** | -0.194 (0.1028) |
| tau TX | 0.083 (0.4869) | 0.036 (0.7611) | -0.148 (0.2163) | -0.21 (0.0772) |
| tau FA | 0.019 (0.874) | 0.019 (0.8741) | -0.148 (0.2153) | -0.126 (0.2914) |
| pT231-tau TBS | 0.098 (0.4107) | 0.013 (0.9162) | 0.072 (0.5465) | 0.072 (0.546) |
| pT231-tau TX | 0.058 (0.6293) | 0.164 (0.1684) | **0.312 (0.0076)** | **0.232 (0.0496)** |
| pT231-tau FA | -0.121 (0.313) | 0.177 (0.1379) | 0.192 (0.107) | **0.289 (0.0138)** |

rs= Spearman correlation coefficient; sol=soluble; ins=insoluble. TBS, TX and FA represent different sequential fractions. P-values < 0.05 were considered statistically significant in this exploratory analysis. Significant correlations are shown in bold.

### Supplemental Table 9: Comparison of tau measurements in between subject groups

| | AD patients (N=72) | | Controls (N=41) | | |
|---|---|---|---|---|---|
| Variable | N | Median (minimum, maximum) or No. (%) of patients | N | Median (minimum, maximum) or No. (%) of patients | P-value |
| Temporal cortex | | | | | |
| sol total tau | 72 | 107743.8 (40783.6, 227510.3) | 41 | 113608.5 (27590.5, 212176.9) | 0.7974 |
| ins total tau | 72 | 38463.0 (2575.7, 155918.1) | 41 | 4929.2 (941.5, 15080.3) | **<0.0001** |
| sol pS396/404-tau | 72 | 830.8 (70.4, 7527.1) | 41 | 144.3 (0.0, 521.3) | **<0.0001** |
| Ins pS396/404-tau | 72 | 68363.6 (196.4, 199058.8) | 41 | 624.3 (0.0, 22494.3) | **<0.0001** |
| Frontal cortex | | | | | |
| sol total tau | 72 | 102324.1 (35991.2, 168711.0) | 41 | 95239.2 (22891.2, 156687.5) | 0.1177 |
| ins total tau | 72 | 45697.8 (1970.9, 124103.5) | 41 | 3538.5 (508.2, 7544.2) | **<0.0001** |
| sol pS396/404-tau | 72 | 334.5 (0.0, 7220.5) | 41 | 78.5 (6.5, 274.5) | **<0.0001** |
| ins pS396/404-tau | 72 | 63434.7 (405.8, 182778.5) | 41 | 246.0 (0.0, 7338.5) | **<0.0001** |

AD=Alzheimer's disease; sol=soluble; ins=insoluble. P-values result from a Wilcoxon rank sum test. P-values P<0.0125 considered statistically significant after applying a Bonferroni correction separately for each cortex. Significant associations are highlighted in bold.

**Supplemental Table 10: UFM1 and UFSP2 protein correlations with DDR gene expression in AD patient temporal cortex**

| | $r_s$ (P-value) | | | |
|---|---|---|---|---|
| DNA damage response gene | sol UFSP2 | ins UFSP2 | sol UFM1 | ins UFM1 |
| TP53BP1 | **0.4427 (0.0003)** | -0.1349 (0.2919) | -0.0556 (0.6652) | **-0.2723 (0.0308)** |
| XRCC4 | -0.0088 (0.9454) | -0.0168 (0.896) | 0.2182 (0.0858) | 0.2089 (0.1004) |
| XRCC5 | **0.3611 (0.0036)** | -0.2241 (0.0775) | 0.1113 (0.3853) | -0.1476 (0.2483) |
| XRCC6 | **0.6963 (<0.0001)** | -0.2271 (0.0735) | 0.1958 (0.1242) | **-0.3413 (0.0062)** |
| PRKDC | 0.1358 (0.2886) | -0.1792 (0.16) | -0.0382 (0.7664) | 0.0029 (0.9819) |
| LIG4 | **0.2484 (-0.0496)** | -0.0724 (0.5726) | 0.061 (0.6346) | -0.2008 (0.1146) |
| DCLRE1C | **-0.5209 (<0.0001)** | 0.1549 (0.2254) | 0.0356 (0.7818) | **0.3793 (0.0022)** |
| NHEJ1 | **-0.4578 (-0.0002)** | -0.0203 (0.8743) | -0.1612 (0.2069) | 0.2408 (0.0573) |
| RAD50 | 0.0406 (0.7518) | -0.0469 (0.7152) | -0.1975 (0.1207) | -0.0274 (0.8312) |
| MRE11A | -0.1941 (0.1274) | 0.0913 (0.4769) | 0.017 (0.8945) | 0.19 (0.1359) |
| NBN | 0.2096 (0.0992) | 0.0495 (0.6999) | **0.2795 (0.0265)** | 0.0218 (0.8651) |
| ATR | **0.5763 (<0.0001)** | **-0.3692 (0.0029)** | -0.0969 (0.45) | **-0.3643 (0.0033)** |
| RPA1 | 0.0605 (0.6377) | 0.0188 (0.8839) | 0.2146 (0.0912) | 0.1928 (0.1301) |
| RPA2 | **0.6216 (<0.0001)** | **-0.3274 (0.0088)** | -0.059 (0.646) | **-0.2874 (0.0224)** |
| RPA3 | **0.5648 (<0.0001)** | -0.2412 (0.0568) | 0.0403 (0.754) | **-0.3803 (0.0021)** |
| RAD52 | **-0.4823 (<0.0001)** | 0.0441 (0.7316) | **-0.2529 (0.0455)** | **0.2562 (0.0427)** |
| BRCA1 | **-0.4733 (<0.0001)** | 0.0874 (0.4958) | -0.1402 (0.2733) | **0.2819 (0.0252)** |
| BRCA2 | **-0.3655 (0.0032)** | -0.1035 (0.4194) | **-0.2735 (0.0301)** | 0.2264 (0.0744) |
| RAD51 | **-0.3311 (0.008)** | -0.0526 (0.6823) | -0.1921 (0.1314) | **0.2903 (0.021)** |
| PALB2 | -0.0689 (0.5916) | -0.0609 (0.6353) | 0.0129 (0.9201) | 0.0246 (0.8484) |
| CHEK1 | **-0.4362 (0.0004)** | 0.0658 (0.6085) | -0.2144 (0.0915) | 0.2 (0.116) |
| ATM | **-0.6344 (<0.0001)** | 0.1036 (0.4192) | -0.1807 (0.1564) | **0.4192 (0.0006)** |
| CHEK2 | **-0.4996 (<0.0001)** | 0.0999 (0.436) | -0.0818 (0.5238) | **0.3096 (0.0135)** |
| TP53 | **-0.305 (0.0151)** | 0.0879 (0.4935) | 0.1123 (0.3811) | **0.2659 (0.0352)** |
| WEE1 | **-0.3486 (0.0051)** | **0.2603 (0.0393)** | **0.2598 (0.0398)** | **0.3429 (0.0059)** |
| MDM2 | **-0.5028 (<0.0001)** | 0.0817 (0.5243) | -0.0732 (0.5685) | **0.3693 (0.0029)** |
| CDKN1A | **-0.3514 (-0.0047)** | 0.2199 (0.0834) | 0.0345 (0.7884) | 0.23 (0.0697) |
| GADD45A | 0.2332 (0.0659) | 0.0599 (0.6411) | 0.0396 (0.7581) | -0.2018 (0.1127) |
| GADD45B | -0.0573 (0.6556) | 0.1422 (0.2661) | 0.0759 (0.5543) | 0.0263 (0.8378) |
| GADD45G | **-0.3806 (0.0021)** | 0.2275 (0.073) | 0.1538 (0.2289) | 0.237 (0.0614) |
| CDK1 | **-0.2935 (0.0196)** | -0.0273 (0.8321) | -0.1468 (0.2509) | 0.0893 (0.4866) |
| CDK2 | **-0.3861 (0.0018)** | 0.1382 (0.28) | 0.1911 (0.1336) | **0.32 (0.0106)** |

rs=Spearman correlation coefficient; sol=soluble; ins=insoluble. P-values < 0.05 are considered statistically significant. Significant correlations are shown in bold. DDR: DNA damage response.

**Supplemental Table 11: UFM1 and UFSP2 protein correlations with seven UPR genes expression in AD patient temporal cortex**

| | rs (P-value) | | | |
|---|---|---|---|---|
| unfolded protein response gene | sol UFSP2 | ins UFSP2 | sol UFM1 | ins UFM1 |
| HSPA5/Bip | -0.1358 (0.2886) | 0.0436 (0.7345) | 0.0255 (0.8427) | 0.0911 (0.4776) |
| EIF2AK3/PERK | **-0.3025 (0.016)** | 0.1514 (0.2363) | -0.048 (0.7088) | 0.2215 (0.081) |
| ATF4 | **-0.5855 (<0.0001)** | **0.2511 (0.0471)** | -0.0812 (0.5269) | **0.3133 (0.0124)** |
| DDIT3/CHOP | **-0.3001 (0.0169)** | 0.0284 (0.8249) | -0.1445 (0.2584) | 0.1391 (0.2769) |
| ERN1/IRE1α | **-0.2489 (0.0492)** | 0.066 (0.6075) | 0.0803 (0.5314) | **0.2606 (0.0391)** |
| XBP1 | -0.1614 (0.2063) | 0.1596 (0.2116) | 0.0664 (0.605) | 0.1689 (0.1858) |
| ATF6 | **0.2696 (0.0326)** | -0.0294 (0.8189) | 0.2133 (0.0932) | -0.0156 (0.9033) |

rs=Spearman correlation coefficient; sol=soluble; ins=insoluble. P-values < 0.05 are considered statistically significant. Significant correlations are shown in bold. UPR: unfolded protein response.
